# Stress-sensitive inference of task controllability

**DOI:** 10.1101/2020.11.19.390393

**Authors:** Romain Ligneul, Zachary Mainen, Verena Ly, Roshan Cools

## Abstract

Estimating the controllability of the environment enables agents to better predict upcoming events and decide when to engage controlled action selection. How does the human brain estimate controllability? Trial-by-trial analysis of choices, decision times, and neural activity in an explore-and-predict task demonstrate that humans solve this problem by comparing the predictions of an “actor” model with those of a reduced “spectator” model of their environment. Neural BOLD responses within striatal and medial prefrontal areas tracked the instantaneous difference in the prediction errors generated by these two statistical learning models. BOLD activity in the posterior cingulate, temporoparietal, and prefrontal cortices covaried with changes in estimated controllability. Exposure to inescapable stressors biased controllability estimates downward and increased reliance on the spectator model in an anxiety-dependent fashion. Taken together, these findings provide a mechanistic account of controllability inference and its distortion by stress exposure.

## INTRODUCTION

Influential theories suggest that the human brain navigates its environment by building predictive models of the world, which in turn fuel cognitive processes such as directed exploration, goal-directed decisions, and forward planning^1–3^. While these internal models can take diverse mathematical forms, their efficiency always depends on the use of task-relevant and cost-efficient state spaces^4–6^. Most often, these state spaces are state-action spaces in which the actions of the agent actively contribute to the prediction of upcoming events. For example, a driver must take into account the movement of their hands to predict the future position of their car. By contrast, a passenger worried for their safety should ignore their own hands and instead focus on the hands of the driver to anticipate potential hazards.

Determining whether an environment is controllable or not is key to deciding to which extent one’s actions should influence the prediction process since only controllable environments afford causal influence over state transitions. Controllable contexts thus prompt the use of “actor” models including one’s own actions as predictors, whereas uncontrollable contexts prompt the use of simpler “spectator” models linking past and future states of the environment. By gating the causal influence of action selection, controllability likely plays a central role in the engagement of elaborate action selection mechanisms. Supporting this idea, it is well established that prior exposure to controllable contexts promotes proactive and goal-directed strategies in a vzariety of cognitive tasks^7,8^. Conversely, the lack of perceived control over events, especially stressful ones, constitutes a well-established correlate and a potential predictor of prevalent psychiatric disorders involving an increased influence of reactive and habitual behaviours, such as depression, anxiety, post-traumatic stress, or obsessive-compulsive disorders^9–13^.

Numerous studies have shown that exposure to uncontrollable stressors can induce a state of learned helplessness characterized by the generalization of passive reactions to subsequent challenges^7,8^. Evidence indicates that this maladaptive state largely depends on functional changes within the medial prefrontal cortex (mPFC) and the serotonin system^14–18^. In humans, a handful of neuroimaging experiments have further suggested that the anterior insula and cingulate cortex contribute to the detrimental effects of uncontrollable stressors^19,20^. Beyond stress induction studies, the sense of being in control of one’s own actions and their outcomes is known to modulate hemodynamic responses parietal and prefrontal cortices^21–23^ and the right temporoparietal junction (TPJ) was found to track the divergence of action-outcome transitions, a feature of controllable environments^24^.

Yet, little is known about the algorithms by which the brain estimates dynamically to what degree a task is controllable. A general strategy is to estimate controllability by computing the causal effect the agent’s own actions have over the environment. Formally, a task can thus be deemed controllable when the transfer entropy (TE) —a generalization of Granger causality to non-linear and discrete systems— linking state and action time-series is positive^25,26^. By comparing the entropy of observed states given previous states and actions [H(S’|S,A)] to the entropy of observed states given previous states only [H(S’|S)], this information-theoretic quantity isolates the effective causal influence of actions over state transitions (**Figure 1a, Supplementary Note 1**). In the vocabulary of causal mediation analysis^27^, positive TE values entail the existence of a natural direct effect linking actions to future states of the environment (**Supplementary Note 2**). Here, we develop a computational model that tracks a dynamic approximation of TE and we use it to shed light on the cognitive and neural mechanisms supporting the ability to infer task controllability and adapt behaviour accordingly.

**Figure 1.**
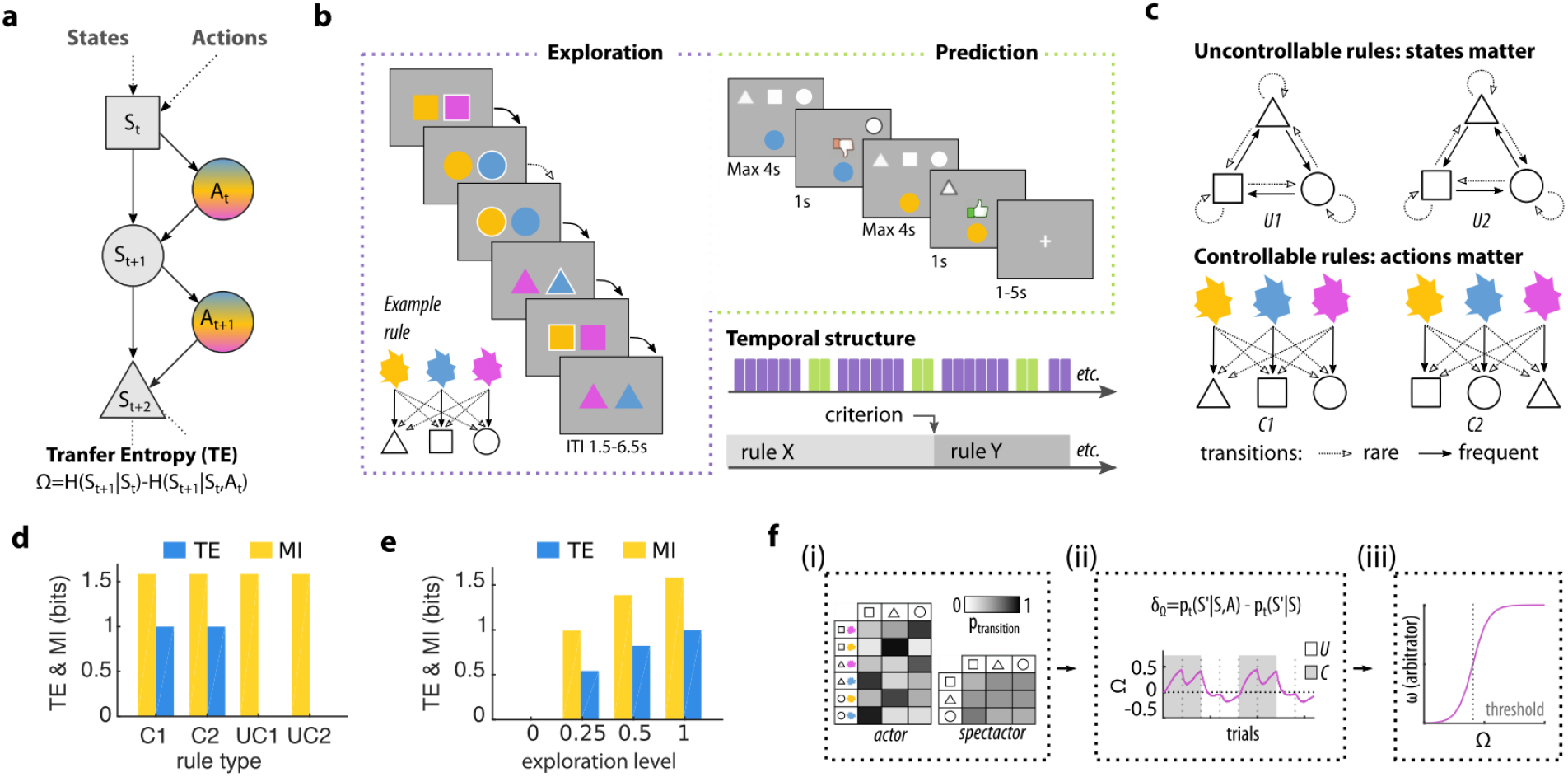
Theoretical framework and experimental protocol. **a**, Controllability can be inferred from transfer entropy, an information-theoretic measure quantifying to which extent a time series causally influences another one. **b**, Time course of a novel explore-and-predict task divided in short mini-blocks. Each mini-block consists of a series of exploratory trials (violet) followed by two counterfactual prediction trials (green) used to assess learning and subjective controllability. **c**, Representation of the 2 uncontrollable rules (U1, U2) and the 2 controllable (C1, C2) rules, which alternate covertly to govern the evolution of the environment. Note that rule C2 was the only rule allowing state repetition, a feature that was taken into account in our analyses. **d**, Simulations showing the dissociation of controllability, as indexed by TE, and predictability, as indexed by the mutual information (MI) shared between successive state-action pairs (random exploration policy). **e**, Simulations under controllable rule C2 showing that TE requires exploration to be used as a proxy for controllability. In the complete absence of exploration, the conditional entropies H(S’|S) and H(S’|S,A) are both null because the agent maintain itself indefinitely in a single preferred state (see also **Supplementary Figure 1C**). **f**, Synthetic overview of the algorithm able to derive an online approximation of TE (termed Ω) by comparing on each trial the transition probabilities of an actor (SAS’) and a spectator (SS’) model of the world. By thresholding Ω, the algorithm could in turn arbitrate between spectator and actor models when making predictions depending on current controllability estimates (ω). This architecture was compared to other controllability estimation schemes and a standard model-based learning model tracking only SAS’ transitions.

Based on this information-theoretic formalism, we designed an explore-and-predict task that allowed us to manipulate controllability and assess the resulting changes in terms of subjective controllability and prediction accuracy. This new task was first used in behavioural (n=50) and fMRI (n=32) experiments which aimed at: (a) demonstrating that humans infer task controllability by estimating an approximation of transfer entropy; (b) establishing the dissociation of spectator and actor models predicted by the transfer entropy hypothesis at the behavioural and neural levels; and (c) unraveling the neural substrates underlying the representation of controllability itself and its influences on behaviour. In a subsequent stress experiment (n=54), we exposed participants to either uncontrollable or controllable electric shocks before administering the explore-and-predict task in order to (d) provide causal evidence supporting a dissociation of the spectator and actor models and (e) test whether learned helplessness can be characterized by an increased reliance on the former relative to the latter.

## RESULTS

### Experimental paradigm and computational model

Healthy human participants were invited to explore an abstract environment composed of three states (square, circle, triangle) and three actions (yellow, blue, magenta). A hidden transition rule always determined upcoming states, either dependent on the action of the participants (controllable rules, C) or the previous state only (uncontrollable rules, U) (**Figure 1b**). The transition rules were probabilistic and reversed covertly, so participants needed to explore and accumulate evidence in order to tell which rule was operative. From time to time, the participant was asked to predict the most likely upcoming state given a state-action pair (e.g. “blue” action in “circle” state), and its counterfactual (e.g. “yellow” action in “circle” state). This procedure yielded a direct yet implicit assessment of their subjective sense of controllability because counterfactual predictions should only differ in controllable contexts, where selected actions determine upcoming states (**Figure 1c**). A novel and distinguishing feature of our task is that controllability varied independently of uncertainty (**Figure 1d**), a methodological improvement over earlier paradigms where the two constructs covary systematically^13,24,28^. Another key difference from previous studies is that we did not include any reinforcers: participants were merely instructed to explore their environment to perform accurate predictions when asked to. Here, controllability estimation can interact with but does not depend on reward and punishment processing^7,29,30^, the only requirement being to maintain a minimal level of exploration, or noise in the action selection process (**Figure 1e, Supplementary Note 1**).

To untangle the mechanisms of controllability estimation in this task, we designed a computational architecture for dynamically tracking an approximation of TE (for a detailed description, see **Supplementary Figure 1a, Methods**). Paralleling the standard computation of TE, two sets of transition probabilities were monitored, one corresponding to an “actor” model (tracking state-action-state transition, SAS’) and the other to a “spectator” model (tracking state-state transition, SS’, **Figure 1f,i**). Following each transition, an approximation of TE (hereafter termed Ω) was updated in proportion to the difference between ‘actor’ and ‘spectator’ transition probabilities pSAS’-pSS’ (**Figure 1f,ii**). Intuitively, this difference term can be understood as an instantaneous causality signal, reflecting how likely the last state transition towards S’ was due to the influence of action A rather than state S. By integrating pSAS’-pSS’ over time, Ω thus reflects the causal influence of actions on recent state transitions (**Supplementary Figure 1b**).

This causality signal Ω is at the core of the proposed algorithm, which arbitrates between the actor and the spectator model when making predictions about upcoming states. Specifically, the relative weight of the actor versus spectator model is set by an arbitrator (hereafter termed ω) whose value can be interpreted as an estimate of controllability. Two parameters influence the mapping between Ω and ω: a threshold determining how much causal evidence is required to infer controllability and a slope determining how fast controllability estimates change around that threshold (**Figure 1f,iii**). This SAS’-SS’-Ω algorithm was contrasted with a conventional model-based learning algorithm^2,31^ and with two models estimating controllability based on the uncertainty —as indexed by the conditional entropy H(S’|A,S)— or divergence of SAS’ transition probabilities bound to different actions (as indexed by the Jensen-Shannon divergence). Importantly, this simpler algorithm could still learn transition probabilities from both uncontrollable and controllable conditions in stable environments, but the lack of controllability-dependent arbitration makes it less efficient in volatile environments alternating rapidly between controllable and uncontrollable rules.

### Controllability drives learning and predictive decisions

Participants performed well on the task: in all experiments, the average prediction accuracy was substantially above chance (**Figure 2a**). In the fMRI experiment, for which participants received more training, accuracy was also stable across conditions and time (**Supplementary Table 1**). Prediction accuracy dropped and then rapidly recovered after covert reversals in transition rules: it already exceeded chance levels on the first pair of prediction trials after reversal (**Figure 2b**). Prediction accuracy also correlated positively with working memory capacity as indexed by d-primes in a standard 2-back task (**Figure 2c**), consistent with the engagement of a model-based learning process^32^.

**Figure 2.**
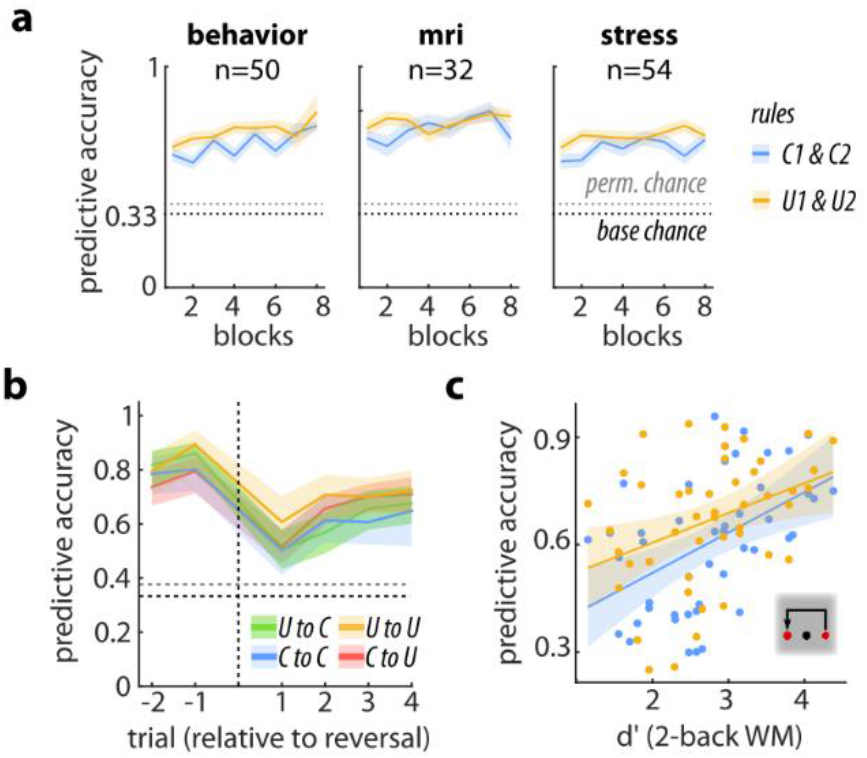
Behavioral performance. **a**, Accuracy in the prediction trials was above chance for both conditions in each of the experiment (behavioural: n=50, t(49)=11.31, p=2.9×10^−15^, d=1.60, 95% CI=(0.20,0.29), fMRI: n=32, t(31)=12.67, p=8.5×10^−14^, d=2.24, 95% CI=(0.24,0.33), stress: n=54, t(49)=11.64, p=3.1×10^−16^, d=1, 95% CI=(0.20,0.28); see also Supplementary Table 1). **b**, Rapid recovery of predictive accuracy for all reversal types (first pair of predictions after reversal: n=50, t(49)=6.64, p=2.4×10^−8^, d=0.94, 95% CI=(0.11,0.20)). Accuracies were split by reversal type for visual purpose (U denotes uncontrollable rules and C denotes controllable rules). **c**, Positive correlation between working memory capacity indexed by a 2-back task (see Supplementary Methods) and predictive accuracy in the explore-and-predict task for controllable (blue: n=46, ρ=0.52, p=1.9×10^−4^, 95% CI (0.18,0.67)) and uncontrollable (orange, n=46, ρ=0.40, p=0.006, 95% CI (0.13,0.62)) contexts. Shaded areas represent standard errors of the mean (SEM).

In line with our prediction that humans solve the task by estimating Ω, Bayesian model comparisons demonstrated that SAS’-SS’-Ω schemes outperformed the conventional model-based learning algorithm (SAS’ alone) in all experiments (**Figure 3a**; **Supplementary Figure 2a**). Simulation analyses confirmed that the model was identifiable and that most of its parameters could be recovered accurately (**Supplementary Figure 2b-d**). As expected, the arbitrator ω captured quantitative changes in subjective controllability, indexed by the proneness of participants to predict that different actions would lead to different states in counterfactual prediction trials (**Figure 3b**). Critically, the SAS’-SS’-Ω scheme which included an arbitration mechanism accounted better for the dynamics of subjective controllability changes around reversals than did the SAS’ model alone (**Figure 3c**, correct prediction of subjective controllability: 72.1% versus 66.5%, N=50, z(49)=3.90, d=0.64, p=9.7×10^−5^, 95% CI (0.004,0.028)). As compared to other controllability estimation schemes (i.e. SAS’-SS’-H and SAS’-SS’-JS), its arbitrator variable ω also predicted variations in subjective controllability more accurately (**Supplementary Figure 2e-f**). The benefits of monitoring controllability are further illustrated by the finding that the likelihood of using the SAS’-SS’-Ω scheme over the SAS scheme increased with accuracy across subjects (**Supplementary Figure 2g**).

**Figure 3.**
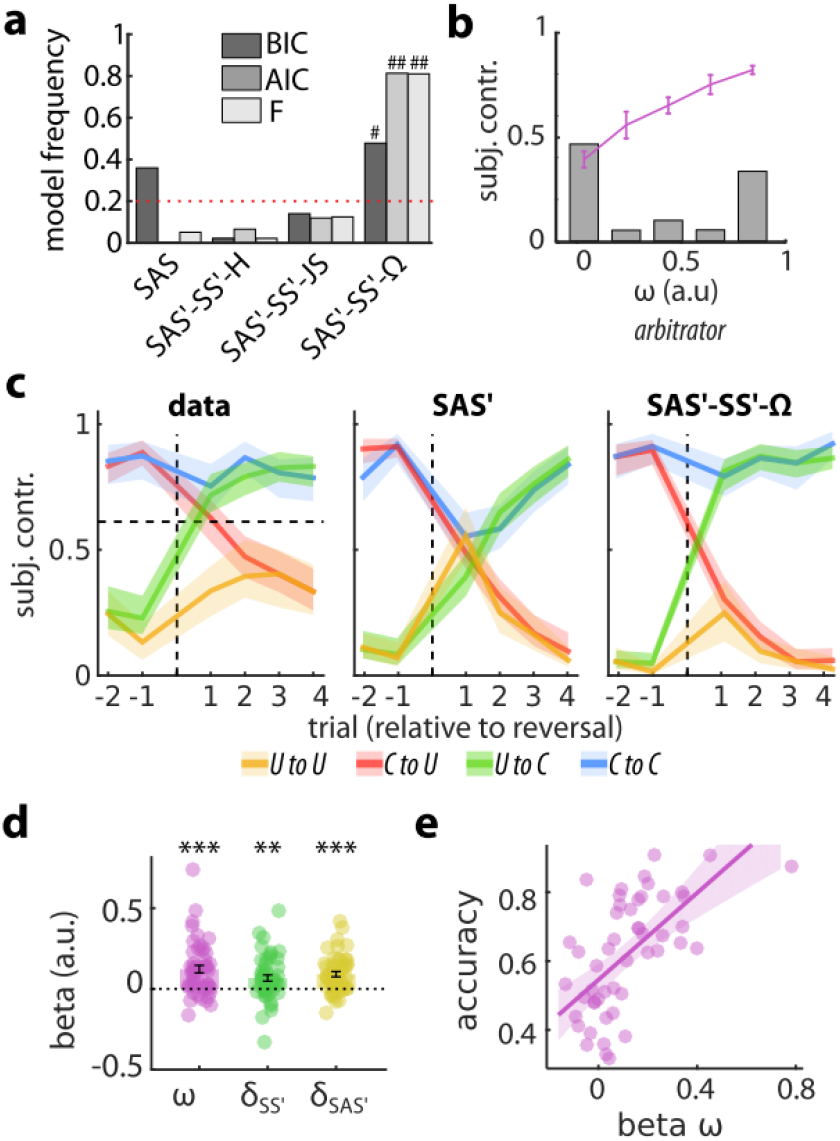
Computational modeling and decision times. **a**, Bayesian group model comparison pooled over three experiments showed the advantage of the model estimating controllability using an approximation of TE (SS’-SAS’-Ω) over the actor model (SAS’) alone and two models estimating controllability based on the uncertainty (SS’-SAS’-H) or the Jensen-Shannon divergence (SS’-SAS’-H) of SAS’ transitions (see Methods and Supplementary Figure 1-2). **b**, Normalized distribution of the arbitrator variable ω (grey bar) and its linear relationship with subjective controllability (pink line). Pairs of prediction trials were labelled as “subjectively controllable” when counterfactual predictions differed (e.g. different responses for blue and yellow actions in the circle state). **c**, The SS’-SAS’-Ω scheme captured the dynamics of subjective controllability around reversals better than the SAS’ model. **d**, Coefficients of the regression predicting log-transformed reaction times in the behavioural experiment using actor and spectator prediction errors as well as ω (N=50; δ_SAS’_: t(49)=6.17, p=1.1×10^−6^, d=0.79, CI (0.6,0.12); δ_SS’_: t(49)=4.39, p=0.001, d=0.49, CI (0.3,0.11); ω: z=4.10, p=4.1×10^−5^, d=0.74, CI (0.08,0.18)). **f**, Positive correlation between the effect of ω on reaction times and predictive accuracy across participants (N=50, ρ=0.67, p=1.1×10^−7^, CI (0.45,0.81)). Error bars and shaded areas represent SEM. ***p<0.001. ^###^p_exceedance_ > 0.999, ^#^p_exceedance_ > 0.95. BIC: Bayesian Information Criterion. AIC: Akaike Information Criterion. F: Free Energy.

Importantly, since only one controllable rule (and none of the uncontrollable rules) allowed immediate state repetitions (rule C2), state repetition events provided a salient psychological cue that contributed to controllability detection. Accordingly, rule C2 was associated with a higher frequency of subjective controllability responses than rule C1 across the three experiments (behaviour: N=50, 82.3+/−13.3% *vs* 75.6+/−14.0%, t(49)=4.1, p=1.5×10^−4^, d=0.58, 95% CI (0.03,0.10); fMRI: N=32, 88.7+/−14.5% *vs* 69.3+/−15.4%; t(31)=6.29, p=5.5×10^−7^, d=0.55, 95% CI (0.13,0.26); stress: N=27, t(53)=4.03, p=1.8×10^−4^, d=0.55, 95% CI (0.04,0.13)). On average, participants also chose more frequently the actions which could lead to state repetition whenever rule C2 was active, thereby indicating that they leveraged this feature of our task to refine controllability inferences in all experiments (behavioural: 53+/−8%, t(49)=2.6, p=0.012, d=0.37, 95% CI (0.007,0.05); fMRI: 51.5+/−4.6% t(31)=1.85, p=0.07, d=0.33, 95% CI (−0.001,0.03); stress: 53.2+/−10.3%, t(53)=2.29, p=0.025, d=0.31, 95% CI (0.004,0.06)). Computational models took this factor into account by allowing prior knowledge about transition rules — as derived from the instruction phase— to constrain the update of actor and spectator models (see Supplementary Information, modelling subsection for more details).

### Dissociation of actor and spectator models

Model comparison results are consistent with our proposal that subjects estimate the subjective controllability of an environment by separately tracking and comparing an actor and a spectator model. In order to further test the dissociation of the actor and spectator models, we used subject-level GLMs to assess trial-by-trial fluctuations of decision times. It is known that decision times slow down following state prediction errors^33,34^. The large amount of exploratory trials per participant thus allowed us to analyze decision times as a proxy of model updating and to evaluate to which extent controllability *per se* influences the speed of action selection [behaviour: 562+/−163 trials; fmri: 550+/−115; stress: 519+/−84]. To do so, we extracted the prediction errors derived from both the actor and spectator models (hereafter termed δ_SAS’_ and δ_SS’_, average correlation: 0.22+/−0.17). We found that both types of prediction errors slowed responding (**Figure 3d**) and independently explained variance in decision times (**Supplementary Figure 3**). We also observed that in periods of higher estimated environmental controllability (i.e. higher ω), decision times were slower. This controllability-dependent slowing correlated positively with predictive accuracy (**Figure 3e**), suggesting that learning in controllable contexts is supported by a more controlled action selection process even when no reinforcement is at stake.

Separable neural correlates should therefore exist for the prediction errors generated by the actor and spectator probability tracking processes, δ_SAS’_ and δ_SS’_. A conjunction analysis first revealed that both types of prediction errors activated the typical set of bilateral brain areas commonly associated with state prediction errors^2,31^, such as the frontoparietal network and the pre-supplementary motor area (**Figure 4a, Supplementary Table 2**). Testing directly the effect of δ_SS’_-δ_SAS’_ (mathematically equivalent to pSAS’-pSS’) using a conventional parametric analysis at the whole brain level showed that the mPFC and the nucleus accumbens encoded negatively this signal required for the update of controllability. In both cases, ROI analyses indicated a positive response to δ_SAS’_ and an absence of a relationship with δ_SS’_ (**Figure 4b-c, Supplementary Table 3**). A similar pattern was observed in the right TPJ (p^FWE^=0.09) and in the dopaminergic nuclei of the brainstem at a more lenient threshold (**Supplementary Figure 4c**). In order to ascertain this dissociation, we performed two additional analyses which fully circumvented the collinearity issues which might arise due to the correlation of prediction errors (average r=0.56+/−0.057). First, we contrasted events where only δ_SS’_ or only δ_SAS’_ were above their respective 66th percentile. Second, we contrasted the parametric effects of δ_SS’_ and δ_SAS’_ derived from two separate first-level GLMs. These analyses confirmed that the NAcc and the mPFC significantly dissociated the two prediction error terms, although only the mPFC systematically survived correction for multiple comparisons (**Supplementary Figure S4a-b, Supplementary Table 3**, for robustness checks).

**Figure 4.**
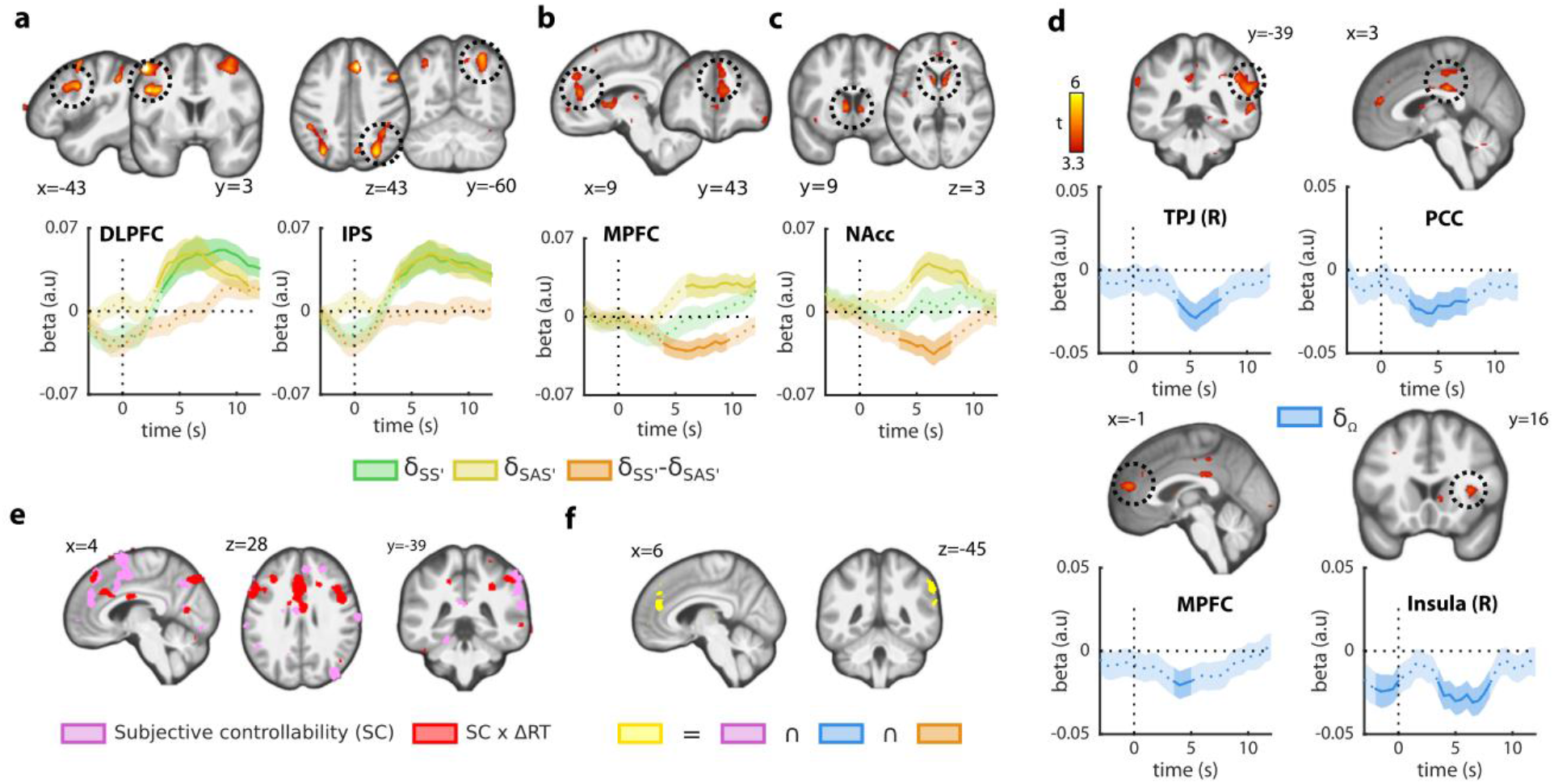
Neuroimaging. **a**, A conjunction analysis revealed brain regions whose activity encoded both δ_SS’_ and δ_SAS’_ positively. **b-c**, Parametric analysis of BOLD responses showed that the mPFC and the nucleus accumbens encoded negatively the difference term δ_SS’_ −δ_SAS’_ used to update controllability. Both regions encoded δ_SAS’_ positively, but showed no clear-cut modulation by δ_SS’_. At a more lenient voxel-wise threshold (p<0.005^UNC^), the right TPJ also survived correction for multiple comparisons (see Table S3 and Figure 4e). **d**, Brain regions encoding signed the second-order prediction errors δ_Ω_. All areas surviving correction for multiple comparisons showed a negative effect, implying greater activity when actions appeared less causal than expected. **e**, Decoding of subjective controllability from brain data. A searchlight analysis based on the six exploratory trials preceding a prediction pair revealed that the mPFC, the posterior dmPFC, the right TPJ, and the precuneus were sensitive to upcoming reports of controllability (pink). Decodability extended to the dlPFC and ACC in participants who displayed slower reaction times in controllable contexts (red). **f**, Spatial overlap (yellow) between the decodability of subjective controllability (4e), controllability prediction errors (4d), and the difference term δ_SS’_ −δ_SAS’_ (4b-c). The right TPJ and the mPFC were the only regions highlighted by each of these analyses (threshold of each map: p^UNC^<0.005). The time courses are shown below (A-D) were only used for robustness checks and visualization. Statistical inferences were based on whole-brain effects at standard thresholds (voxel-wise: p<0.001, uncorrected; cluster-wise: p<0.05^FWE^). Shaded areas represent SEM.

### Neural correlates of dynamic controllability

Having established the dissociation of δ_SAS’_ and δ_SS’_ at the behavioural and neural levels, we next probed the correlates of the prediction error δ_Ω_ governing changes in estimated controllability (δ_Ω_=δSS’-δ_SAS’_-Ω_t-1_). This second-order learning mechanism is key to accumulate, over time, evidence in favor or against the controllability of the ongoing rule. Whole-brain analyses revealed a significant negative relationship between δ_Ω_ and neurovascular responses in the posterior cingulate (PCC), the right dorsal anterior insula (dAI), the right temporoparietal junction (TPJ), and the mPFC (**Figure 4d, Supplementary Table 4**). Mixed-effects ROI analyses including decision times and Ω confirmed that these effects reflected a genuine response to controllability prediction error, peaking 4-8 seconds after trial onset.

In order to unravel the neural correlates of controllability with maximal sensitivity, we performed a multivoxel pattern analysis (MVPA). A support vector machine classifier was trained at predicting whether streaks of consecutive exploratory trials would result in an implicit report of subjective controllability or not in the upcoming pair of prediction trials (i.e. different or identical responses for each counterfactual). Whole-brain maps of classification accuracy were obtained using the searchlight method (leave-one-run-out cross-validation). Local patterns of activity in the precuneus, the right TPJ, the supplementary motor area (SMA), the left premotor cortex, the left dlPFC (**Figure 4e, Supplementary Table 5**) contained information relative to subjective controllability. Interestingly, the decoding performance in the ACC and the right dlPFC scaled with the lengthening of decision times in controllable contexts from one participant to another. The dorsal bank of the mPFC and the right TPJ were the only two regions whose activity was simultaneously sensitive to subjective controllability (as probed by MVPA), to controllability prediction errors and to the instantaneous difference between δ_SAS’_ and δ_SS’_ (**Figure 4f**).

### Uncontrollable stressors promote the spectator model

We applied this new paradigm to better understand the computational mechanisms underlying learned helplessness. More precisely, we hypothesized that exposure to uncontrollable stressors might bias controllability estimation mechanisms to promote reliance on the spectator model relative to the actor model. We invited participants to perform an active avoidance task exposing them to mild electric shocks before completing the explore-and-predict task (**Figure 5a**). Participants in the controllable group learned to avoid the shock following one of the three possible cues by pressing the correct response button (out of six alternatives). Shocks received by participants in the uncontrollable group were yoked to the former so that their decisions did not influence shock probability. As expected, this procedure induced a dissociation between actual shock frequency, matched across groups by design, and reported shock expectancy (**Figure 5b**), so that shock expectancy remained high until the end of the induction phase in the uncontrollable group.

**Figure 5.**
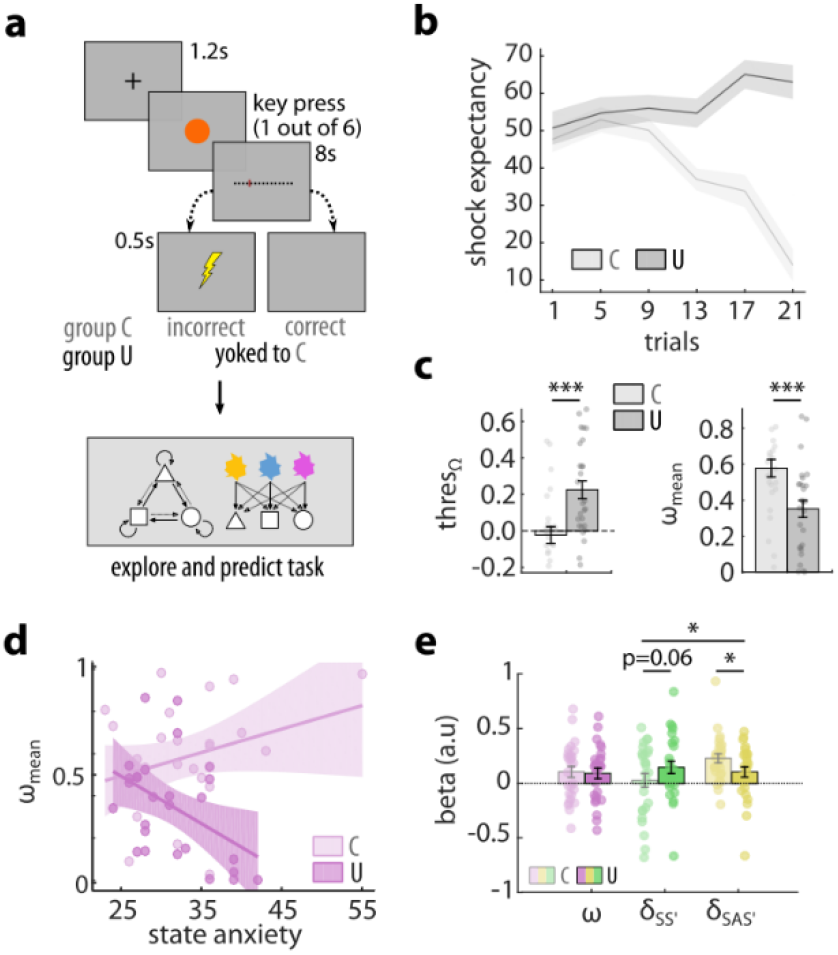
Stress experiment. **a**, Induction of controllable and yoked uncontrollable stress followed by the explore-and-predict task. **b**, Temporal evolution shock expectancy during the induction phase, split by condition. **c**, Impact of induction type on the parameter *thres*_Ω_ and the average value of the arbitrator ω. The value above which Ω was treated as evidence for a controllable environment increased significantly following uncontrollable stressors (N=27 per group, t(52)=4.56, p=2.7×10^−4^, d=1.1, 95% CI (0.120.39)), resulting in increased reliance on the spectator model when making predictions, as indexed by the reduction of ω at the group level (N=27 per group, z=2.60, p=0.009, d=0.77, 95% CI (0.08,0.29)) see also Supplementary Figure S5 and Table S6). **d**, State anxiety moderated the effect of induction type on the arbitrator variable (ω) reflecting controllability estimation. Higher state anxiety was associated with greater reliance on the spectator model after exposure to uncontrollable stressors (dark pink, N=27, r=-0.46, p=0.015, 95% CI (−0.70,-0.08)) but not after controllable stressors (bright pink, N=27, r=0.30, p=0.12, CI (−0.16,0.61)). Correlation coefficients were significantly different between uncontrollable and controllable group (z=2.87, p=0.002). **e**, The impact of induction type on the slowing of decision times induced by δ_SS’_, δ_SAS’_ (average correlation of the two first-order prediction errors: 0.45+/−0.17) was consistent with this increased reliance on the spectator model (interaction group by PE type: F(1,52)=5.97, p=0.018, η^2^=0.07, 95% CI (0.005,0.20)). All error bars and shaded areas represent SEM. *p<0.05,**p<0.01.

Despite the absence of aversive reinforcers in the explore-and-predict task, we observed clear carry-over effects from the shock experiment when analyzing the model parameters governing controllability estimation (**Figure 5c, Supplementary Table 6**). In particular, the threshold parameter increased significantly in the uncontrollable group compared with the controllable group (**Figure 5c**, left). This parameter determines how much causal evidence is required before controllability is inferred. Therefore, when making predictions, participants exposed to uncontrollable stressors relied more on the spectator model, demonstrated by the reduced average value of the arbitrator ω (**Figure 5c**, right) as well as the direct analysis of subjective controllability estimates, revealing that counterfactual predictions were more often identical in the uncontrollable group (z=1.69, p=0.045, d=0.48, 95% CI (0.006,0.10), one-tailed).

We found no statistically significant evidence that the stress manipulation affected the overall motivation of ability to perform the task. Accuracies (U=0.61+/−0.16; C=0.65+/−0.16%; t(52)=0.81, p=0.43, d=0.22, 95% CI (−0.05,0.13)) and decision times (U=2.07+/−0.73s; C=1.97+/−0.65s; t(52)=0.56, p=0.59, d=0.14, 95% CI (−0.48,0.27)) did not differ significantly across groups in prediction trials. Furthermore, when restricting our analysis to the uncontrollable group, the SAS’-SS’-Ω model still outperformed a variety of simpler models, including the standard SAS’ model (actor only), a SS’ model (spectator only), and a RL model which only learned through the feedback delivered on predictive trials (**Supplementary Figure 5a**). Exposure to uncontrollable stressors thus elicits an imbalance between actor and spectator mechanisms for transition probability learning consistent with a sustained shift in controllability expectations. This finding provides a parsimonious account of the cross-contextual generalization of passive strategies, a core feature of helpless states^7,8^. Interestingly, state anxiety, as assessed before the experiment, moderated the induction of controllability estimation biases. It predicted the average value of the arbitrator only in participants exposed to uncontrollable stressors (**Figure 5d**).

Since uncontrollable stressors promoted increased reliance on the spectator model and decreased reliance on the actor model, we expected PE-dependent slowing effects to follow a similar pattern. Confirming this prediction, the type of stress induction profoundly altered the slowing of decision times by actor and spectator prediction errors (**Figure 5e**). Post-hoc tests showed that the effect of δ_SAS’_ was significantly lower (t(52)=-2.45, p=0.018, d=0.65, 95% CI (−0.13,-0.1)) and that the impact of δ_SS’_ was marginally higher (δ_SS’_: t(52)=1.86, p=0.068, d=0.51, 95% CI (−0.006,0.16)) in the uncontrollable group compared with the controllable group (**Figure 5e**).

## DISCUSSION

Taken together, these findings shed light on one of the most fundamental aspects of human experience: the ability to estimate to which extent our actions affect our environment and to adjust our decisions accordingly. Our results demonstrate that this ability involves the comparison of actor and spectator models of the ongoing task, which are dissociable computationally, behaviourally and neurally. In turn, controllability estimates can be used to arbitrate between these models when making predictions about future events. The mPFC and the striatum encode the difference between the prediction errors generated by each model, while signals related to the update of controllability estimates are found in a more posterior brain network encompassing the TPJ and the PCC. Furthermore, exposure to uncontrollable stressors biases this process assessed by the explore-and-predict task, hence establishing its relevance for the study of psychiatric disorders involving altered perceptions of controllability ^9,12,13,35^.

Historically, the concept of task controllability has been heavily influenced by learned helplessness studies in which animals granted the ability to actively terminate stressors are compared to yoked animals exposed to the exact sequence of stressors, but whose actions are made independent from stressor termination^8,14,15^. In this line of research, focused on the long-lasting consequences of stress exposure, more controllable contexts were defined as those in which the mutual information linking the timings of actions and stressor offsets is higher^36^. However, a positive mutual information linking an organism’s actions and upcoming states of the environment is a necessary but not sufficient condition to declare a task controllable. For example, the highly positive mutual information linking the statements of a weather forecaster with the occurrence of rain should obviously not be interpreted as a sign that the forecaster controls the weather because the statements of the forecaster and the occurrence of rain are both conditioned by past meteorological states. Moreover, following this incomplete definition, variations of task controllability were often obtained by manipulating uncertainty about future states^17,20,28^, hence leading to ambiguous conclusions regarding the mechanisms underlying the estimation of controllability *per se* and its downstream influence on behaviour.

Formalizing controllability using transfer entropy (TE) rather than mutual information allowed us to design a task in which controllability varied independently from uncertainty. In addition, this approach provided an algorithm for detecting genuine changes in task controllability. Model comparisons showed that, across the three experiments, algorithms monitoring controllability using an approximation of TE (i.e SAS’-SS’-Ω scheme) accounted better for participants’ choices than a simpler SAS’ learning algorithm. While the neural correlates of state prediction errors elicited by the explore-and-predict task were remarkably similar to those reported in two-step decision tasks analyzed using SAS’ (or actor) learning^2,37,38^, model comparisons revealed the concomitant engagement of SS’ (or spectator) learning. Crucially, the SAS’-SS’-Ω model also outperformed two architectures in which controllability was derived from statistical features of the SAS’ transition probabilities only: namely, they exploited the uncertainty (SAS’-SS’-H) or the divergence of counterfactual SAS’ transitions (SAS’-SS’-JS) to discriminate between controllable and uncontrollable contexts. While they predicted qualitatively similar behaviours (e.g. dynamics of subjective controllability change around reversals), these alternative models were less efficient at capturing the dynamic fluctuations of subjective controllability during the task. Finally, the analysis of reaction times confirmed that participants were sensitive to the prediction errors generated by the spectator and actor models, whose comparison governed the update of controllability estimates in the SAS’-SS’-Ω scheme only.

The actor and the spectator can be viewed as two state-spaces competing to structure the learning of statistical contingencies. When controllability estimates are low, the spectator model representing only the successive states of the environment dominates. In contrast, when controllability estimates are high, the actor model representing both states and actions takes over. By defining the most appropriate state space dynamically, controllability estimation improves predictions about the future states of one’s environment and can therefore contribute to maximizing utility when reward or punishment rates depend on such predictions. And by promoting reliance on a simpler spectator model when the environment is deemed uncontrollable, it can also minimize the metabolic cost and subjective effort associated with controlled action selection^39,40^. These hypotheses could be tested directly by introducing reinforcers in the explore-and-predict task, but it is already worth noting that the controllability-dependent arbitration logics can readily explain why Pavlovian (equivalent to SS’) and instrumental (equivalent to SAS’) learning mechanisms are respectively promoted in uncontrollable and controllable contexts^28^. Furthermore, the preferential encoding of actor prediction errors by the mPFC, the striatum, and the dopaminergic midbrain is consistent with earlier findings showing that the mesolimbic pathway preferentially encodes reward prediction errors in instrumental learning tasks^6,41–43^. The finding that the actor and spectator models only dissociated in these deep structures close to the midline is consistent with a recent MEG study showing that the human brain exploits shared “neural codes” to address sensorimotor and perceptual demands in controllable and uncontrollable contexts^44^.

By comparing the predictions emanating from the actor and spectator models, one can derive an instantaneous causality signal (i.e how likely did the last action cause the last state transition). Encoded by mPFC and striatal BOLD responses, this instantaneous signal can then be integrated over time, hence reflecting the causal influence of actions over recent transitions. A signature of the second-order prediction errors supporting this integration was found in the right TPJ, the dorsal mPFC, the right insula, and the PCC. The right TPJ and the mPFC were the only regions sensitive to the difference of the two first-order prediction errors, to these second-order prediction errors used to update controllability, as well as to subjective controllability as assessed by the decoding analysis. They are thus strong candidates for the implementation of controllability monitoring in our task. Supporting this view, the right —but not left— TPJ has previously been found to encode the divergence in action-outcome distributions^24^ and the discrepancy between expected and actual outcome timings in a simple sensorimotor task alternating controllable and uncontrollable trials^23^. It is also consistent with a study showing that mPFC lesions can alter the perception of controllability in simple instrumental learning tasks^45^. Given that the mPFC and the uncertainty of SAS’ transitions are involved in the trade-off between model-based and model-free decision-making systems^46^, these results support the emerging hypothesis^7,47^ that perceived or expected controllability may play a role in the relative influence of these systems as assessed by two-step tasks^31,48^.

Other prefrontal areas were sensitive to variations in subjective controllability according to the decoding analysis, likely reflecting adaptations of brain networks to task controllability^49^ or the contribution to controllability estimation of cognitive processes which were not captured by the SAS’-SS’-Ω scheme. For example, the ACC and the anterior insula —which encoded controllability prediction errors— play an established role in the signaling of state uncertainty or task volatility, which may both participate in controllability estimation^28,50,51^. Strikingly, a higher sensitivity of the dlPFC and dmPFC (extending ventrally to the dorsal ACC) to controllability was also associated with a stronger influence of controllability on decision times. This finding suggests that controllability detection may foster a form of proactive response inhibition previously linked with dlPFC and dmPFC activity^52,53^. The engagement of more elaborate action selection processes may ultimately depend on the valuation of control itself, which is known to involve dorsal ACC activity^54^. Indeed, while this process is usually studied by varying task difficulty, it is clear that, by gating the causal influence of actions, variations of task controllability moderate the expected benefits of exerting cognitive control.

Having described the computational principles and outlined neural correlates of controllability estimation, we sought to test whether an experimental manipulation could alter this process and simultaneously contribute to a better understanding of the learned helplessness phenomenon. Indeed, exposure to uncontrollable stressors is known to induce passive responses to subsequent controllable stressors, but the origins of this maladaptive strategy remain poorly understood. In particular, it is unclear whether prior exposure to uncontrollable stressors induces an increased sensitivity to future aversive events, reduces the expectation of control with respect to future stressors, or reduces expectations of control in general^8,29^. Our results support the latter hypothesis by showing sustained alterations of controllability estimation in human participants previously exposed to uncontrollable versus controllable stressors. More precisely, the specific increase observed for the threshold parameter implies that the former group needed to integrate more causal evidence before considering a given rule as controllable in the explore-and-predict task. The dorsal anterior insula (dAI) is involved in the modulation of pain perception by controllability^20^ and it was found to encode controllability prediction errors in our fMRI experiment. Therefore, prior exposure to uncontrollable stressors may have altered dAI excitability to distort subsequent controllability estimation mechanisms. Supporting this idea, a study showed that a lower perception of control mediates the exacerbation of dAI responses to physical threats in more anxious individuals, who also displayed lower controllability estimates following uncontrollable stressors in our data^19^. Yet, this increased reliance on the spectator relative to the actor model following uncontrollable stressors likely involves several other brain areas, including the mPFC and the dorsal raphe nucleus, both highly sensitive to stressor controllability^14–16^.

In sum, the explore-and-predict task allowed us to isolate the core computations supporting the inference of task controllability by excluding reinforcers and matching uncertainty across contexts. The mPFC and the right TPJ emerged as the two most promising candidates for the neural implementation of controllability inference and it would therefore be interesting to confirm their causal contribution using brain stimulation methods in the future. By showing that the human brain can compute an approximation of transfer entropy, our study may help to bridge the gap between neuroscience and artificial intelligence research, where transfer entropy plays an important role in solving unsupervised learning problems^55,56^. Investigating in greater detail the interactions between controllability estimation and model-based reinforcement learning mechanisms will constitute an important step in this direction. More invasive techniques will also be required to understand how these computations are implemented within local neural circuits and how neuromodulators such as dopamine or serotonin mediate their broad impact on stress responses and mental health^7,14,15,57^.

## METHODS

### Participants

For the behavioural experiment, fifty young adult participants (mean age: 24.7, range: 18—43, 27 women) were recruited via the Sona system (human subject pool management system) of the Radboud University (The Netherlands). All participants were included in the data analysis. For the fMRI experiment, thirty-two young adult participants (mean age: 25.1, range: 20—43, 18 women) were recruited through the same system. For the stress experiment, a total of 62 participants (mean age = 21.8; range: 18-27, 52 women) were recruited via the Sona system of Leiden University. One additional participant was excluded a posteriori from the fMRI experiment and four participants were excluded from the stress experiments, together with their yoked counterparts (see Supplementary Methods for details on exclusion and inclusion criteria). The behavioural and fMRI experiments were approved by the local ethics committee (CMO region Arnhem/Nijmegen, The Netherlands, CMO2001/095). The stress experiment was approved by the Psychology Research Ethics Committee (CEP17-0905/282) at Leiden University. All participants provided written informed consent, in line with the declaration of Helsinki and were compensated for their participation in the study (10€/hour for the behavioural and fMRI experiments, 7.5€/hour for the stress experiment).

### Explore-and-predict task

In the 3 experiments, the overall structure of the task was identical. Participants performed 6 (fMRI and stress experiment) or 7 (behavioural experiment) exploratory trials before a pair of predictions were required. Pairs of predictions always probed the two actions available for a given state (e.g blue followed by yellow in the circle state), to derive subjective controllability from counterfactual responses. Participants received feedback about their predictions in 50 percent (fMRI and stress) or 100 percent (behavioural experiment) of the trails. In the fMRI and stress tasks, feedback was delivered only after one of the two counterfactual predictions in order to prevent participants from inferring whether the rule was controllable or not based on feedback.

On each exploratory trial, two identical geometrical shapes were displayed side by side. The color of each shape determined the action corresponding to left and right button presses (side randomly assigned in each trial). An urgency signal was displayed after 1.5s. Transitions to the next state were always governed by one of the four rules, as displayed in Figure 1c. In order to maximize the variation of prediction errors, the transitions were stochastic (noise: 0.05 to 0.2).

The first prediction trial of each pair was simply displayed at the end of the ITI of the previous exploratory trial. An urgency signal was displayed after 4s. The hypothetical state action pair was displayed at the centre of the screen, just below a question mark, and the 3 possible next states were displayed as white geometrical shapes at the top of the screen. The selected state was then highlighted and the feedback was displayed when applicable.

The ongoing rule was never changed before 4 pairs of predictions were completed. In the behavioural experiment, the rule changed from then as soon as 5 correct responses were provided in the last 6 predictions or if the last 4 predictions were accurate. In the fMRI experiment, the rule was changed as soon as the p-value of a binomial test indicated that accuracy was significantly below chance (p<0.05, one-tailed, chance level: 1/3), hence making the accuracy threshold more lenient as the number of predictions made for a given rule increased. In all experiments, the rule changed after 10 pairs of predictions, even if performance did not meet the learning criterion. Prediction trials were pseudo-randomly ordered with the constraint that each state would be tested a similar number of times. An exhaustive description of instructions, counterbalancing, reversal schedules, transition noises, and timings is available in Supplementary Methods.

### Stress induction task

To test the impact of prior controllability over stress on subsequent controllability estimations, participants underwent a stressor controllability manipulation before the explore-and prediction task in a between-subjects design. Critically, we employed a between-subjects yoked control procedure to match the amount and order of aversive outcome stimuli between the controllable and uncontrollable conditions. We randomized participants in blocks of four where the controllable condition of a yoked pair was always administered first in order to create the schedule for the yoked counterpart in the uncontrollable condition. Electric stimuli served as stressors in the manipulation task and were delivered by a Digitimer DS7 stimulator. First, individual levels of intensity of the electric stimulus for the manipulation task were determined using a stepwise procedure in which the intensity of the stimulus was gradually increased until participants reported a ‘just bearable, but not yet painful’ experience of shock. A yoked control-design with pre-programmed pseudorandomized schedule enabled us to match the amount and order of electric stimuli between the conditions.

In the controllable condition, a total of four cues (different in shape and color) were presented for at least six repetitions each following a pre-programmed pseudorandomized schedule. Participants could learn by trial-and-error the correct response corresponding to the cue (a key between 1 and 6) to avoid the electric stimulus. Critical trials on which participants would be able to prevent the electric stimulus for the first time according to the schedule were repeated until the participants arrived at a correct response. As such, all participants underwent the whole schedule with a minimum of 24 trials and were able to acquire the correct response for each cue.

The uncontrollable condition was yoked to the controllable condition, such that participants experienced a comparable pattern of events across conditions. However, in the uncontrollable condition, participants were not able to acquire these action-outcome contingencies to prevent the shocks, whose sequences were merely replayed from the yoked participants performing the controllable condition. Data collection and analysis were not performed blind to the conditions of the experiments. An exhaustive description of counterbalancing, instructions and procedures is available in Supplementary Methods.

### Computational modelling

The main purpose of all SAS’-SS’-Ω algorithms is to provide a way to dynamically estimate the causal influence of actions over state transitions by updating a variable termed Ω. In all models, S represents the previous state of the environment, A represents the previous action and S’ represents the current state of the environment. The local causality estimate Ω can only be used as a proxy for controllability, which is not a property of actions but of the environment. It is this “inferred controllability” variable, termed ω, which can then be used to decide (arbitrate) whether one should make predictions using learned S-S’ transitions or learned SA-S’ transitions. Ω is homologous to transfer entropy (TE, which is itself a generalization of Granger causality to discrete and non-linear domains). See Supplementary Methods for a detailed explanation of the differences between TE and Ω.

In order to demonstrate that participants used a dynamic estimate of transfer entropy to solve the task, we compared the SAS’-SS’-Ω algorithm to a standard model-based architecture tracking SAS’ transitions^2^. This latter algorithm corresponds to the actor model alone. Its asymptotic performance in stable environments is identical to that of SAS’-SS’-Ω algorithms. We also compared the SAS’-SS’-Ω algorithm to alternative controllability estimation schemes based solely on statistical features of SAS’ transitions: namely, the uncertainty of SAS’ transitions (SAS’-SS’-H, lower during periods of high controllability) and the Jensen-Shannon divergence of counterfactual SAS’ transitions (SAS’-SS’-JS, higher during periods of high controllability). These two alternatives schemes are fully described in Supplementary Information.

The actor model tracks transitions linking state-action pairs to newly encountered states (i.e. SAS’). It updates transition probabilities in the following fashion.

Realized transitions:

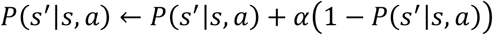

Unrealized transitions:

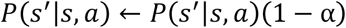

Where α ∈ [0,1] controls to which extent learned transition probabilities are determined by the most recent transitions. The prediction error 1−P(s’| s,a) is noted δ_SAS’_ in the main text.

The spectator model tracks transitions linking states to newly encountered states (i.e. SS’). Therefore, it updates transition probabilities exactly like the actor model, except that only states are represented: P(s’| s,a) is simply replaced by P(s’| s) in the two equations above, and the prediction error 1−P(s’| s) is noted δ_SS’_ in the main text.

Following the update of the actor and spectator models, we allowed prior transition probabilities derived from the instruction phase to constrain the update of each model. This was done by multiplying (element-wise) the relevant vector of probabilities by the corresponding vector of prior probabilities. For example, after a transition from a blue circle (state 2, action 2), the transition probabilities of the spectator model were multiplied by [0.5 λ 0.5] (reflecting the fact that states did not repeat under uncontrollable rules) and those of the actor model were multiplied by [λ 0.5 0.5] (reflecting the fact that the triangle state never appeared after choosing blue under uncontrollable rules). Thus, for any λ<0.5, this prior injection step constrained the update of the spectator and actor model in a way that reflected the transitions *a priori* possible under uncontrollable and controllable rules, respectively. By altering the prediction errors elicited by each model, prior injection had an indirect influence on controllability estimation. For example, a lower value of λ leads to a greater increase of estimated controllability following state repetition events, by reducing the prediction error generated by the actor model relative to the spectator model (see below).

The variable Ω supports the controllability estimation process by tracking the expected difference P(s’| s,a) − P(s’| s) dynamically (or, equivalently, δ_SS’_ − δ_SAS’_). The logic of this process is that, in a controllable environment, actions contribute to predicting the upcoming states and therefore P(s’| s,a) > P(s’| s). Higher values of Ω, therefore, imply higher evidence that the environment is controllable. The update of Ω is governed by the following equation:

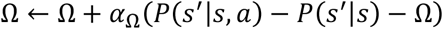

Where αΩ ∈ [0,1] is the learning rate controlling to which extent Ω is determined by the most recent observations.

Since Ω reflects the causal influence of one’s action over state transition, it can be used as a proxy to infer whether the environment is likely controllable or uncontrollable. In order to form the arbitration term reflecting this inference and accommodate inter-individual differences at this step, Ω is thus transformed using a parametrized sigmoïd function:

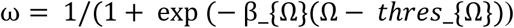

Where thres_Ω_ ∈ [−1,1] corresponds to the threshold above which Ω is interpreted as evidence that the environment is controllable and where β_Ω_ ∈ [0,Inf] determines to which extent evidence that the environment is controllable (i.e. Ω − thresΩ > 0) favors reliance on learned SAS’ transitions when making predictions (and vice-versa for SS’ transitions when Ω−thres_Ω_ < 0). Thus, the variable ω implements the arbitration between the “actor” and the “spectator” model.

When only SAS’ learning is considered, the probability that a given state S’=i will be observed given S and A is directly given by:

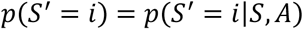

When the SS’-SAS’-Ω architecture is used, the probability that a given state S’=i will be observed given S, A, and ω is directly given by:

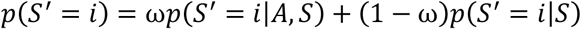

In turn, the probability that the participant predicts the next state would be i (e.g. a square) when confronted to the hypothetical state-action pair S,A (e.g. circle state, blue action) is given by:

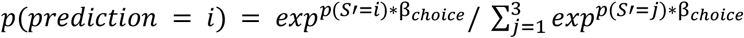

Where β_choice_ ∈ [−Inf,Inf] determines to which extent the participants will systematically select the most likely transition (i.e. the highest p(S’=i), according to what has been learned) to make their predictions. A very positive β_choice_ implies that the participant systematically selects this most likely transition. A β_choice_ around 0 implies that the participant mostly makes random guesses. And a β_choice_ very negative would imply that the participant mostly goes against what he/she has learned.

The full model space was composed of SAS’ alone (2 parameters), SAS’-SS’-Ω (6 parameters), SAS’-SS’-H (5 parameters), and SAS’-SS’-JS (5 parameters). An exhaustive description of these models is available in Supplementary Information.

### Model fitting procedures

Model fitting was performed using a Variational Bayesian (VB) estimation procedure using the well-validated VBA toolbox^58^. The fitting procedure only attempted to explain decisions made in prediction trials. In other words, the decisions made in exploratory trials only indirectly constrained the fit by determining the information gleaned between pairs of prediction trials. For the behavioural experiments, the prior distributions of the various learning rates and the threshold parameter were innately defined as Gaussian distributions of mean 0 and variance 3, which approximates the uniform distribution over the interval of interest after sigmoid transformations. The prior distributions of βchoice and βω parameters were defined as Gaussian distribution of mean 0 and variance 10. For the fMRI and the stress experiments, the prior distributions of every parameter were defined using the posterior mean and variance obtained from the 50 participants who passed the behavioural experiment. Hidden states corresponding to transition probabilities were systematically initialized at 1/3 (equiprobability prior), while Ω was initialized at 0. Detailed information about parameter transformation, model fitting, model comparison, and simulation procedures is available in Supplementary Information.

### fMRI: acquisition

All images were collected using a 3T Siemens Magnetom Prismafit MRI scanner (Erlangen, Germany) with a 32-channel head coil. A T2*-weighted multiband echo-planar imaging sequence with acceleration factor 8 (MB8) was used to acquire BOLD-fMRI whole-brain covered images (TR = 700 ms, TE = 39 ms, flip angle = 52, voxel size = 2.4 × 2.4 × 2.4 mm3, slice gap = 0 mm, and FOV = 210 mm). This state-of-the-art sequencing protocol was optimized from the recommended imaging guidelines of the Human Connectome Project, with the fast acquisition speed facilitating the detection and removal of non-neuronal contributions to BOLD changes (protocols.humanconnectome.org/HCP/3T/imaging-protocols). The experiment was divided into 4 blocks lasting on average 7.7+/−2.1 minutes (662+/−179 volumes). We recorded participants’ heartbeats using the scanner’s built-in photoplethysmograph, placed on the right index finger. Respiration was measured with a pneumatic belt positioned at the level of the abdomen. Anatomical images were acquired using a T1-weighted MPRAGE sequence, using a GRAPPA acceleration factor of 2 (TR = 2300ms, TE = 3.03 ms, voxel size = 1×1×1mm, 192 transversal slices, 8° flip angle). Field magnitude and phase maps were also acquired.

### fMRI: preprocessing

fMRI data processing and statistical analyses were performed using statistical parametric mapping (SPM12; Wellcome Trust Centre for Neuroimaging, London, UK). For each session, the first 4 volumes were automatically discarded by the scanner. Functional images were slice-time corrected, unwarped using the field maps, and realigned to the mean functional image using a rigid-body registration. Functional images were then coregistered to the anatomical T1. Next, the anatomical images were segmented based on tissue prior probability maps for spatial normalisation using the DARTEL algorithm, and the resulting normalization matrix was applied to all functional images. Finally, all images were spatially smoothed with a 6mm Gaussian kernel, except in the decoding analysis for which unsmoothed images were used.

### fMRI analyses

Statistical analyses of fMRI signals were performed using a conventional two-levels random-effects approach in SPM12. All general linear models (GLM) described below included the 6 unconvolved motion parameters from the realignment step. We also included the eigenvariate of signals from cerebrospinal fluid (CSF) in our GLM (fourth and lateral ventricular). Moreover, we used a retrospective image correction (RETROICOR) method to regress out physiological noise, using 10 cardiac phase regressors and 10 respiratory phase regressors obtained by expanding cosines and sines of each signal phase to the 5th order. We also included time-shifted cardiac rates (lag: +6, +10 and +12s) and respiratory volume (−1 and +5s) as nuisance regressors. All regressors of interest were convolved with the canonical hemodynamic response function (HRF). All GLM models included a high-pass filter to remove low-frequency artifacts from the data (cut-off = 96s) as well as a run-specific intercept. Temporal autocorrelation was modeled using an AR(1) process. All motor responses recorded were modeled using a zero-duration Dirac function. We used standard voxel-wise threshold to generate SPM maps (p<0.001 uncorrected), unless notified otherwise. All statistical inferences based on whole-brain analyses satisfied the standard multiple comparison threshold (p(FWE)<0.05) at the cluster level unless notified otherwise. The cluster-size correction was based on random field theory. Prediction error and (log-transformed) decision time regressors were systematically z-scored within individual blocks to exclude scaling effects. All GLM models included separate onset regressors for motor responses, for prediction trials, and for the first trial of each exploratory sequence (where no prediction error was elicited). All models also included parametric regressors for reaction time and ω (reflecting controllability estimates) on prediction trials. A detailed description of the GLMs used to analyze neuroimaging data is available in Supplementary Methods. These GLMs only differ in the way exploratory trials were treated.

In order to verify the robustness of our whole-brain results and inspect the time course of our parametric effects of interest, we performed mixed-effects analyses on BOLD signals filtered and adjusted for nuisance regressors. This adjusted signal was extracted from the functional clusters uncovered by whole-brain analyses and segmented into trial epochs from −3 to +16 seconds around the onset of each exploration trial (excluding the first of each streak). We then estimated the effect of each regressor of interest, at each time point, for all subjects simultaneously. Subject identity was included as a random effect and a subject-specific intercept was included. Parametric regressors were z-scored in the same way as in the mass univariate analyses. Importantly, this approach was not used for statistical inference — since doing so would constitute double-dipping — but merely for visualization purposes. Decoding analyses were performed using the TDT toolbox^59^. Each mini-block of 6 exploratory trials was arbitrarily coded as +1 (controllable) or −1 (uncontrollable) based on the responses given in the upcoming prediction pair (identical responses = uncontrollable; different responses = controllable). We used a leave-one-run out cross-validation scheme with 100 permutations per subject, so that classes remained balanced for training. The training was performed on the beta values associated with each mini-block using a Support Vector Machine (SVM) classifier (L2-loss function, cost parameter set to 1, Liblinear, version 1.94), without feature selection or feature transformation. Since we did not constrain the testing sets to have balanced classes, balanced accuracies were used when reporting the results of the searchlight analysis (performed within an 8mm sphere) at the whole-brain level.

### Statistical procedures

Model selections relied on Bayesian model comparisons and exceedance probabilities, as implemented by the VBA toolbox^58^. The analysis of predictive accuracies over time and across conditions relied on a 2-way repeated measure ANOVAs or one-sample t-tests, assuming normal distribution of the data following *arcsin* transformation. In order to assess whether predictive performance was significantly superior to chance, we permuted correct responses for each hypothetical state independently and compared these permuted responses with actual predictions (1000 permutation per participants). The resulting empirical chance levels were indeed higher than the theoretical level of ⅓ in all experiments (behavioural: 38.0+/−2.5%; fMRI: 39.4+/−1.6%; stress: 38.6+/−2.6%), reflecting the fact that participants knew that 3 transition rules out of 4 did not allow state repetitions.

The analysis of decision times was performed in two steps: first, a logistic regression was performed on binarized decision times (median-split); second, group-level significance was assessed by means of one-sample t-tests. For the analysis of decision times, we excluded trials in which decision times were 3 standard deviations above the mean. Comparison between conditions relied on paired t-tested and comparison between groups (stress experiments) relied on two-sample t-tests, unless normality assumptions were violated, in which case non-parametric equivalents were used (Wilcoxon signed rank and rank sum tests, respectively). All t-tests were two-sided unless notified otherwise. Correlations were based on Pearson coefficients unless normality assumptions were violated, in which case Spearman rank coefficients were used. Confidence intervals were computed using a bootstrapping approach (2000 permutations).

## DATA AVAILABILITY

The experimental paradigm and the code used to generate the figures is available at the following address: github.com/romainligneul/NHBcontrollability. Second-level SPM images are available at the following address: identifiers.org/neurovault.collection:8810. Anonymized data can be accessed at the following address using ORCID identification: <u>data.donders.ru.nl/collections/di/dccn/DSC_3017049.01_905?5</u>.

## CODE AVAILABILITY

The scripts used to collect and analyse data is available upon publication at the following address github.com/romainligneul/NHBcontrollability.

## ACKNOWLEDGEMENTS

We are grateful to Michael Frank for his constructive comments on the manuscript and computational models. We thank Paul Gaalman for his help with fMRI data acquisition. We thank Korina Elefteriadou, Fili Dianellou, Kristin Koelbel, and Julia Breen and SOLO labsupport Leiden University for their help and support in the acquisition of the data for the stress experiment.

## Funding

This work was supported by grants from the Fyssen Foundation and the behaviour and Brain Research foundation awarded to RL (Young Investigator 2017) and a Vici award from the Netherlands Organisation for Scientific Research to RC (NWO 453-14-005). The funders had no role in study design, data collection and analysis, decision to publish or preparation of the manuscript.

## AUTHOR CONTRIBUTIONS

Conceptualization: RL, RC, VL. Methodology: RL, VL. Software & formal analysis: RL. Investigation: RL, VL. Resources: RC, ZM. Data Curation: RL, VL. Writing - Original Draft: RL. Writing - Review & Editing: RL, ZM, VL, RC. Visualization: RL. Funding acquisition: RL, ZM, VL, RC.

## COMPETING INTERESTS

The authors declare no competing interests.

## Notes

### Competing Interest Statement

The authors have declared no competing interest.

### Summary of Updates

Final version

