## Supplementary Material for "Stress-sensitive inference of task controllability"

### 1 Participants

#### 1.1 Behavioural experiment

Fifty young adult participants (mean age: 24.7, range: 18—43, 27 women) were recruited via the Sona system (human subject pool management system) of the Radboud University (The Netherlands). Participants were prescreened to ensure normal to corrected vision, a sufficient understanding of English language, no history of neurological of psychiatric diseases. Pregnancy and use of psychoactive drugs in the 2 weeks prior to the experiments were part of the exclusion criteria. All participants were included in the data analysis. They were compensated at a fixed rate of 8 euros/hour for their participation in the study. The study was approved by the local ethics committee (CMO region Arnhem/Nijmegen, The Netherlands, CMO2001/095) and all participants provided written informed consent, in line with the declaration of Helsinki.

#### 1.2 fMRI experiment

Thirty-two young adult participants (mean age: 25.1, range: 20—43, 18 women) were recruited via the human subject pool management system of the Radboud University (The Netherlands). Participants were pre-screened and excluded using the same criteria than for the behavioural experiment. One participant was excluded *a posteriori* from the study due to excessive sleepiness in the scanner (eyes-closed more than 10 % of the time n=1). All participants were compensated at a fixed rate of 10 euros/hour for their participation in the study. Most participants were also included in the behavioural experiment or in an previous pilot study (n=29) but inclusion in the fMRI experiment was independent of performance in the behavioural experiment. The study was approved by the local ethics committee (CMO region Arnhem/Nijmegen, The Netherlands, CMO2001/095) and all participants provided written informed consent, in line with the declaration of Helsinki.

#### 1.3 Stress experiment

A total of 62 participants (mean age = 21.8; range: 18-27, 52 women) were recruited via SONA, the human participants system, at Leiden University. Exclusion criteria were: pregnancy, deviation from normal or corrected-to-normal vision, insufficient understanding of written and spoken English, self-reported history of any neurological, cardiac, or psychiatric disease, any use of prescribed psychotropic drugs within two weeks or alcohol in the 24 hours prior to the experiment. The study was approved by the Psychology Research Ethics Committee (CEP17-0905/282) at Leiden University. All participants provided written informed consent and were compensated for their participation: 7.50 euros/hour or 2 participant credits. Four participants needed to be excluded for different reasons: 1. no aversive stimuli being delivered due to cable being connected to the wrong port; 2. stressor manipulation task crashed during the experiment; 3. unable to finish the experiment due to delays; 4. noncompliance to instructions of the experimenter. To retain the yoked design, we also excluded the four yoked counterparts of these participants. For another four participants in the uncontrollable condition, an error occurred in the yoking procedure resulting in one aversive stimulus less for these participants. Nevertheless, we decided to include these participants and their yoked counterparts in the analyses: if anything, this single extra aversive stimulus in the controllable condition would only make the comparison with the uncontrollable condition more conservative. Thus, after exclusion of 8 participants, a total sample size of 54 was left for analyses.

### 2 Explore-and-predict task

#### 2.1 Instructions and training

Upon arrival on site and after completing the informed consent form, participants were told that they would take part to an experiment probing the human ability to detect, learn and use the rules which govern a simplified environment composed of three states and three actions. The experimenter explained that states corresponded to the geometrical shapes displayed on the screen (triangle, circle, square), whereas the actions related to the colors of these shapes (blue, yellow, magenta). Participants were told that they would have to explore this simplified environment by selecting, in each state, an action using the left/right arrows of the keyboard (behavioural and stress experiments) or button box (fMRI experiment). They were then presented with a scheme, similar to that of Fig. 1c, describing the 4 possible rules which controlled the transitions from one state to another.

The experimenter explained the distinction between controllable and uncontrollable rules and told participants that their goal would be to learn the rule during exploration and that, from time to time, the computer program would ask them to predict the next most likely state given a hypothetical state-action pair. The experimenter explained the logic of counterfactual testing, making sure that participants would understand the implications of making the same prediction for the two possible actions associated with a given state (i.e no decisive influence of actions implying a "spectator" rule); versus making distinct predictions (i.e. decisive influence of actions implying an "actor" rule).

The participants were told that rules would alternate covertly, without warning, and that they should thus constantly evaluate whether the rule they learned was still valid. They were also informed that the number of trials in the task would depend on their ability to make explicit predictions, but the exact criterion determining covert reversals was not revealed.

Following these verbal instructions, participants completed a training phase divided in 4 blocks (behavioural) or 8 blocks (stress and fMRI experiments). At the beginning of each block, the rule which governed the transitions was displayed explicitly on the screen. Participants were told to "feel what it is like to explore the environment when such a rule is active" and they were given the opportunity to make predictions from time to time, as in the testing phase. Positive or negative feedback was delivered following each prediction. Once the participants had completed all training blocks, the experimenter verified that they had understood the principle of the experiment. Next, they explained that, unlike the training phase, the transitions of the test phase would be slightly noisy. This noise was described as a small proportion of trials in which the computer program would select randomly the next state instead of applying the active rule. It was made explicit that, in the prediction trials, the noise would be absent and that the feedback - if any - would faithfully reflect the active rule.

The relatively long duration of the training phase (15-20 minutes) was needed to ensure that most participants would get the principle of the task. A pilot study had suggested that only a subset of participants were able to operate successfully in the task with minimal instructions and training. Note that participants in the fMRI experiment completed the training phase twice, once outside the scanner (4 blocks) and once inside the scanner (4 blocks) during the acquisition of the anatomical scan.

#### 2.2 Structure of the task

In the 3 experiments, the overall structure of the task was identical. Participants performed 6 (fMRI and stress experiment) or 7 (behavioural experiment) exploratory trials before a pair of predictions was required. Pairs of predictions always probed the two actions available for a given state (e.g blue followed by yellow in the circle state), in order to derive subjective controllability from counterfactual responses. Participants received feedback about their predictions in 50 percent (fMRI and stress) or 100 percent (behavioural experiment) of the trails. In the fMRI and stress tasks, feedback was delivered only after one of the two counterfactual predictions in order to prevent participants from inferring whether the rule was controllable or not based on prediction feedback.

On each exploratory trial, two identical geometrical shapes were displayed side by side. The visual angle encompassing both shapes was about 6-7° for all experiments. The color of each shape determined the action corresponding to left and right button presses (the side was randomly assigned in each trial). In the behavioural experiment, these shapes were displayed for at least 0.5s or until the participant made a choice. The shapes started to fade out automatically after 1.1s in the fMRI experiment and the fading was complete after 1.5s (in the behavioural experiment, shapes faded out at button press in 0.25s). In every case, a small warning symbol urging participants to make a choice appeared between the two shapes if no button press had been registered after 1.5s. followed by a blank screen whose duration was:

- Uniformly distributed in the [0.25, 0.5] interval (mean 0.375s) for the behavioural experiment
- Exponentially distributed in the [0.05 5] interval (mean 2s) for the fMRI experiment.
- Exponentially distributed in the [0.05 4.5] interval (mean 0.5s) for the stress experiment.

The first prediction trial of each pair was simply displayed at the end of the ITI of the previous exploratory trial. These trials were self-paced, although a warning appeared on screen after 4s to urge participants to make a choice. The hypothetical state action pair was displayed at the center of the screen (visual angle of about 2°), just below a question mark, and the 3 possible next states were displayed as white geometrical shape at the top of the screen. The selected state was then highlighted for 0.3s. In the next period of 0.7s the feedback was displayed (if no feedback was displayed, the selected state simply remained highlighted during 0.7s). In the behavioural experiment, a blank screen lasting 0.3s followed each prediction trial. In the fMRI and stress experiements, it was followed by a blank screen whose duration was distributed similarly to the ITI describe above (0.05 to 4.5 or 5, respectively).

#### 2.3 Reversal schedule

The ongoing rule was never changed before 4 pairs of predictions were completed. In the behavioural experiment, the rule changed from then as soon as 5 correct responses were provided in the last 6 predictions or if the last 4 predictions were accurate. In the fMRI experiment, the rule was changed as soon as the p-value of a binomial test indicated that accuracy was significantly below chance ($p<0.05$, one-tailed, chance level: 1/3), hence making the accuracy threshold more lenient as the number of predictions made for a given rule increased. In all experiments, the rule changed after 10 pairs of predictions, even if performance did not meet the learning criterion. Inability to reach this criterion could be due to distorted controllability perception or simply a slow, suboptimal learning process. Therefore, We did not exclude any data based on performance. Predictions were pseudo-randomly ordered with the constraint that each state would be tested a similar number of times.

Contrary to the stress and fMRI experiments which were divided in 4 identifiable blocks separated by short pause screen showing a slide describing the possible rules, the version of the SS'SAS' task used for the behavioural experiment was not explicitly divided into blocks. In all experiments, the computer program covertly alternated rules to enable studying changes in predictions following 4 uncontrollable (U) to controllable (C) reversals, 4 C to U reversals, 2 C-C reversals (e.g. from rule C1 to rule C2) and 2 U-U reversals (e.g. U1 to U2). Because inter-block transitions were not taken into considerations (the participants were informed that each block was independent), stress and fMRI experiments thus required 4 blocks in which the 4 rules were tested (3 transitions per blocks). Block types were counterbalanced across participants. The 4 possible blocks were: $\{U1, U2, C2, C1\}$, $\{U2, C1, U1, C2\}$, $\{C1, C2, U2, U1\}$ and $\{C2, U1, C1, U2\}$, hence resulting in 8 cross-dimensional (i.e U to C, C to U) and 4 intra-dimensional (i.e U to U, C to C) reversals. This ordering enabled us to distinguish amongst different computational models and to promote the use of a controllability monitoring strategy. Note that each rule was tested once in each position within blocks.

##### 2.4 Transition noise

Finally, in order to increase the variance and the temporal distribution of prediction errors related to $p(S'|S)$, $P(s'|a,s)$ and $\Omega$, the transitions were noisy. In the behavioural experiment, a random noise level was set to 7.5% (so that the active rule would be applied in 92.5% of the exploratory trials). In the fMRI experiment, the noise level depended on the transition. Within each rule, the three possible SA-S' or S-S' transitions had thus a noise of 5, 10 or 20% (e.g. under U1, square to circle would be realized 95% of the time while circle to triangle would be realized only 80% of the time). This refined procedure was used to increase the variance of the prediction error terms regressed onto neural activity without completely disrupting learning in less efficient participants (the lowest noise transitions being still easily detectable). Finally, in the stress experiment, the noise level was set to 10% for all transitions.

#### 3 Other tasks

##### 3.1 Working memory task

The working memory (WM) task was programmed using Psychtoolbox (Matlab 2014b). It tested four different conditions: 2-back, 3-back, 2-forth and 3-forth. In every condition, participants were presented with a constant stream of numbers (display duration: 700ms, inter-trial interval: 800ms. In the n-back blocks, they were instructed to press the space bar each time a number would be identical to the number encountered 2 or 3 trials before, depending on the difficulty level. In the n-forth task, participants were asked to detect streak of 3 or 4 numbers increasing or decreasing in a row (data not analysed). Each block was composed of 50 trials and included eight targets. Incorrect responses (i.e. miss or false alarm) triggered a warning message (1s). Before the task itself, participants performed 16 trials (2 targets) of each condition. Working memory performance was assessed using a d-prime criterion (z(HIT)-z(FA)). Since the d-prime could not be computed in one participant who had difficulty completing the 3-back task, we restricted our analysis to the 2-back task. Note that 4 more participants in the behavioural experiment did not complete the working memory task due to technical problems, hence resulting in

46 participants included in WM-related analyses.

#### 3.2 Stress induction task

To test the impact of prior controllability over stress on subsequent controllability estimations, participants underwent a stressor controllability manipulation prior to the explore-and-prediction task in a between-subjects design. Critically, we employed a between-subjects yoked control procedure in order to match the amount and order of aversive outcome stimuli between the controllable and uncontrollable conditions. We randomized participants in blocks of four (two Controllable and two Uncontrollable conditions) where the Controllable condition of a yoked pair was always administered first in order to create the schedule for the yoked counterpart in the Uncontrollable condition.

##### 3.2.1 General procedure

Upon arrival to the laboratory, participants were briefly reminded about the experimental procedure. The study was framed such that it was not clear that the study involved a between subject manipulation of stressor controllability but rather that it concerned decision making processes and physiological measurements. After providing informed consent, electrodes for the physiological measurements were positioned on the participant skin conductance, blood pressure, electrocardiogram (ECG) and impedance cardiography (ICG). Subsequently, they filled out several self-report questionnaires (State-Trait Anxiety Inventory, Leiden Index of Depression Sensitivity-Revised, Cognitive Emotional Regulation Questionnaire, Behavioural Activation/Inhibition Scale, Barratts Impulsiveness Scale, Childhood Trauma Questionnaire), and performed a working memory task (2-Back task, see below). However, only the State subscale of the State-Trait Anxiety Inventory was scored and analyzed in the current study.

After a 5 minutes baseline measure of skin conductance, blood pressure, ECG, and ICG while watching a neutral video clip, participants received instructions and training for the explore-and-predict task. Subsequently, they went through a procedure to determine the shock intensity to be used during the stressor controllability manipulation task. By providing the instructions and training for the explore-and-predict task before exposure to any shock stimuli, we prevented any effects of the stressor on the instructions and training of the explore-and-predict task.

Online measures of skin conductance, blood pressure, ECG and ICG were taken during the stressor controllability task (data not analysed). Participants were asked to indicate their levels of positive and negative affect and cognition on a visual analogue scale on different moments during the experiment (baseline, before and after the manipulation, and at the end of the experiment)

##### 3.2.2 Stressor controllability manipulation

Electric stimuli served as stressors in the manipulation task and were delivered by a Digitimer DS7 stimulator. First, individual levels of intensity of the electric stimulus for the manipulation task were determined using a stepwise procedure in which the intensity of the stimulus was gradually increased in intervals of 0.10 mA until participants reported a 'just bearable, but not yet painful' experience of shock on a scale from 0 = not uncomfortable at all to 100 = just bearable, but not yet painful.

Depending on the condition to which participants were randomly assigned (controllable or uncontrollable condition), perceived control over the stressor was manipulated via the presence or absence of objective control (i.e. choice and action-outcome-contingency) respectively. A yoked control-design with preprogrammed pseudorandomized schedule enabled us to match the amount and order of electric stimuli between the conditions as well as to minimize interindividual variation in the manipulation.

In the controllable condition, a total of four cues (different in shape and color) were presented for at least six repetitions each following a preprogrammed pseudorandomized schedule. Participants could – supposedly – learn by trial-and-error the correct response corresponding to the cue (a key between 1 and 6) to avoid the electric stimulus. They were instructed that each cue had a unique corresponding key as correct response and that the correct response would always prevent the electric stimulus. Unbeknownst to the participants, the trial at which they could prevent the electric stimulus for a particular cue for the first time depended on a combination of the preprogrammed schedule and optimal exploration of the participant. For example, participants could prevent the electric stimulus for cue A at the third repetition (A3) for the first time, if they explored a third key on this trial that was different from the first two attempts (for A1 and A2). This third key was then assigned as the correct key corresponding to cue A for the rest of the task. If participants repeated unsuccessful attempts for a specific cue or chose correct keys assigned to other cues, they would receive an electric stimulus. Critical trials on which participants would be able prevent the electric stimulus for the first time according to the schedule were repeated until the participants arrived at a correct response. As such, all participant underwent the whole schedule with a minimum of 24 trials, and were able to acquire the correct response for each cue.

The uncontrollable condition was yoked to the controllable condition, such that participants experienced a comparable pattern of events across conditions. However, in the uncontrollable condition, participants were not able to acquire these action-outcome contingencies to prevent the shocks, but were instead instructed to press a random button (key 1 to 9) on each trial.

### 4 Computational modelling

The main purpose of all SAS'-SS'-Ω variants is to provide a way to dynamically estimate the causal influence of actions over state transitions by updating a variable termed Ω. In all models, S represents the previous state of the environment, A represents the previous action and S' represents the current state of the environment. The local causality estimate Ω can only be used as a proxy for controllability, which is not a property of actions but of the environment. It is this "inferred controllability" variable, termed $\omega$, which can then be used to decide (arbitrate) whether one should make predictions using learned S-S' transitions or learned SA-S' transitions. Ω is homologous to transfer entropy (TE, which is itself a generalization of Granger causality to discrete and non-linear domains), with 3 important differences:

- Ω is not computed based on post-hoc transition probabilities (i.e after all transitions have been observed) but on transition probabilities estimated trial-by-trial (using a delta rule). This means that Ω is dynamic and not static. Unlike TE, Ω can therefore accumulate causal evidence locally in order to discriminate between periods of high controllability and periods of low controllability.

- For the same reason, Ω can take negative values. This can happen just after rule reversals or when the active rule is suddenly violated due to noise. Indeed, in such cases, S-S' transitions can momentarily appear more likely than SA-S' transitions.

- It is not log-transformed and therefore does not represent “bits” of information. Instead, it represents the expected difference between the probability of observations given previous states and actions p(S’lS,A) and the probability of observations given previous states only $(S'|S)$. Importantly, by forcing $p(S'|S,A) \geq (S'|S)$ and by tracking the expected value of $log(p(S'|S,A)) - log(p(S'|S))$ instead of $p(S'|S,A) - p(S'|S)$, $\Omega$ and TE converge towards the same values in the long-run.

In order to demonstrate that participants used a dynamic estimate of transfer entropy to solve the task, we systematically compared variants of the SAS’-SS’-Ω architecture to a standard model-based architecture tracking SA-S’ transitions [1,2]. This approach is powerful because the asymptotic performance of this latter model is identical to that of SAS’-SS’-Ω models, whose relative advantage lies in the ability to arbitrate between controllable (i.e SAS’) and uncontrollable (i.e S-S’) transition matrices to perform predictions. Indeed, the actor (ie. SAS’) model can perfectly learn the state-state (SS’) transition probabilities tracked by the spectator model. It only needs more data points, especially when the number of possible state-action pairs is high. This remark is important because it implies that comparing the SAS’-SS’-Ω architecture to the SAS’ model alone constitutes a fair and stringent test.

In the following, the computational steps relevant for the entire model space are described. While the models took into account all trials providing information about transition probabilities to update hidden states, the fitting procedure only attempted to explain decisions made in prediction trials. In other words, the decisions made in exploratory trials did not constrain the values of the best-fitting parameters.

#### 4.1 Update of SAS’ transitions ("actor model")

This module tracks SA-S’ transitions. Because actions are explicitly represented, it is called the "actor" module. In each exploratory trial and prediction trial followed by a feedback, this standard model-based learning module updates the transition probabilities linking one state-action pair to the newly encountered state in the following fashion:

Realized transitions

$$P(s'|a,s) \leftarrow P(s'|a,s) + \alpha_{sas'}(1 - P(s'|a,s))$$

Unrealized transitions

$$P(s'|a,s) \leftarrow P(s'|a,s)(1-\alpha_{sas'})$$

Where $\alpha_{sas'} \in [0,1]$ controls to which extent learned transition probabilities are determined by the most recent transitions. Note that $1-P(s'|a,s)$ is noted $\delta_{SAS'}$ in the main text.

In order to allow prior knowledge about possible task conditions to constrain the update of the actor model, P(s'la,s) was multiplied element-wise by a vector representing prior transition probabilities such that:

$$P(s'|a,s) = P(s'|a,s) \cdot P_{prior}(s'|a,s) / \sum P(s'|a,s) \cdot P_{prior}(s'|a,s)$$

with $P_{prior}(s'|s)$ being equal to:

| | S1 | S2 | S3 |
|---|---|---|---|
| A1 | 0.5 | $\lambda$ | 0.5 |
| A2 | $\lambda$ | 0.5 | 0.5 |
| A3 | 0.5 | 0.5 | $\lambda$ |

For $\lambda = 0$, the table above represents the transition probabilities which could be expected under the two controllable rules before exploration (i.e. based on task instructions). For $\lambda < 0.5$, *prior injection* had thus a beneficial influence on learning. For $\lambda < 0.5$ (a situation which happened rarely), *prior injection* had on the contrary a detrimental influence on learning. In Fig. S1a, $w_{priorSAS'} = P_{prior}(s'|a,s) / \sum P(s'|a,s) \cdot P_{prior}(s'|a,s)$.

#### 4.2 Update of SS' transitions ("spectator model")

This module tracks S-S' transitions. Because actions are not represented, it is called the "spectator" module. In each exploratory trial and prediction trial followed by a feedback, this spectator module updates the transition probabilities linking one state to the newly encountered state in the following fashion:

Realized transitions

$$P(s'|s) \leftarrow P(s'|s) + \alpha_{ss'}(1-P(s'|s))$$

Unrealized transitions

$$P(s'|s) \leftarrow P(s'|s)(1-\alpha_{ss'})$$

Where $\alpha_{ss'} \in [0,1]$ controls to which extent learned transition probabilities are determined by the most recent transitions. Note that $1 - P(s'|a,s)$ is noted $\delta_{SS'}$ in the main text.

In order to allow prior knowledge about possible task conditions to constrain the update of the spectator model, P(s'ls) was multiplied element-wise by a vector representing prior transition probabilities such that:

$$P(s'|s) = P(s'|s) \cdot P_{prior}(s'|s) / \sum P(s'|s) \cdot P_{prior}(s'|s)$$

with $P_{prior}(s'|s)$ being equal to:

| | S1 | S2 | S3 |
|---|---|---|---|
| S1 | $\lambda$ | 0.5 | 0.5 |
| S2 | 0.5 | $\lambda$ | 0.5 |
| S3 | 0.5 | 0.5 | $\lambda$ |

For $\lambda = 0$, the table above represents the transition probabilities which could be expected under the two uncontrollable rules before exploration (i.e. based on task instructions). For $\lambda < 0.5$, the *prior injection* step thus reduces the probability of state repetitions according to the spectator model. In Fig. S1a, $w_{prior} = P_{prior}(s'|s) / \sum P(s'|s) \cdot P_{priorSS'}(s'|s)$.

#### 4.3 Update of $\Omega$

This second-order module tracks the expected difference $P(s'|a,s) - P(s'|s)$ dynamically (or, equivalently, $\delta_{SS'} - \delta_{SAS'}$). The logic of this process is that, in a controllable environment, actions contribute to predicting the upcoming states and therefore $P(s'|a,s) > P(s'|s)$. At any given moment, a positive $\Omega$ therefore constitutes evidence that the environment is controllable:

$$\Omega \leftarrow \Omega + \alpha_\Omega (P(s'|a,s) - P(s'|s) - \Omega)$$

or, equivalently:

$$\Omega \leftarrow \Omega + \alpha_\Omega (\delta_{SS'} - \delta_{SAS'} - \Omega)$$

Where $\alpha_\Omega \in [0,1]$ is the learning rate controlling to which extent $\Omega$ is determined by the most recent observations.

#### 4.4 Inference of controllability ($\omega$) from $\Omega$

As written above, $\Omega$ reflects the causal influence of one's action over state transition. It can therefore be used as a proxy to infer whether the environment is likely controllable or uncontrollable. In order to form the arbitration term reflecting this inference and accommodate inter individual differences at this step, $\Omega$ is thus transformed using a parametrized sigmoïd function:

$$\omega = \frac{1}{1+\exp(-\beta_{\Omega}(\Omega - threshold_{\Omega}))}$$

Where $threshold_{\Omega}) \in [-1,1]$ corresponds to the threshold above which $\Omega$ is interpreted as evidence that the environment is controllable and where $\beta_{\Omega} \in [0, Inf]$ determines to which extent evidence that the environment is controllable (i.e. $\Omega - threshold_{\Omega} > 0$) favors reliance on learned SAS' transitions when making predictions (and vice-versa for SS' transitions when $\Omega - threshold_{\Omega} < 0$).

In other words, the variable $\omega$ implements the arbitration between the "actor" and the "spectator" model.

#### 4.5 Inference of controllability based on H(S'|A,S)

As an alternative to $\Omega$-based controllability estimation scheme, the agent might in principle derive controllability at the time of predictions based on the uncertainty bound to learned transition probabilities from the actor model. Indeed, in period of high controllability, this uncertainty is on average lower than in period of low controllability. In this case, the arbitrator $\omega$ is given by:

$$\omega = \frac{1}{1+\exp(-\beta_{H}(1.585 - H(S'|A,S) - threshold_{H}))}$$

with:

$$H(S'|A,S) = \sum_{i=1}^{3} -p(S'=i|A,S)\log_2(p(S'=i|A,S))$$

The above expression is greatly simplified as compared to the standard formulation of conditional entropy because A and S correspond to the hypothetical states and actions bound to the ongoing prediction trial, rather than to the entire space of possible starting states and actions. Note also that H(S'|A,S) takes a minimum value of 0 for deterministic transitions (maximal control) whereas it takes a value of 1.585 when transition probabilities are uniform (i.e., $p(S'=i|A,S) = 1/3$ for all i, no control).

##### 4.6 Inference of controllability based on Jensen-Shannon divergence

As a second alternative to Ω-based controllability estimation scheme, the agent might in principle derive controllability from the Jensen-Shannon (JS) divergence of transition probabilities predicted by the actor model. Indeed, JS divergence can quantify to which extent state transition probabilities differ depending on which action is chosen. In this case, the arbitrator $\omega$ is given by:

$$\omega = \frac{1}{1 + \exp(-\beta_{JS}(JS - threshold_{JS}))}$$

with:

$$JS = 0.5 * \sum_{j=1}^{2} \sum_{i=1}^{3} p(S' = i|C = j, S)(\log_2(p(S' = i|C = j, S)) - M)$$

and:

$$M = \log_2(0.5 * \sum_{j=1}^{2} \sum_{i=1}^{3} p(S' = i|C = j, S))$$

Note that in the above expression M merely corresponds to a normalization factor ensuring that $JS \in [0, 1]$ and that $C = \{1, 2\}$ corresponds to the two counterfactual actions available in a given state (e.g. blue and yellow in the circle state). Thus, the more dissimilar the transition probabilities associated with C=1 and C=2, the higher the JS divergence: JS=1 for fully dissimilar distributions (e.g. [0 0 1] and [1 0 0], maximal control) and JS=0 for fully similar distributions (e.g. [0 0 1] and [0 0 1], no control).

##### 4.7 Prediction

When only SAS' learning is considered, the probability that a given state S'=i will be observed given S and A is directly given by:

$$p(S' = i) = p(S' = i|S, A)$$

When the SS'-SAS'-Ω architecture is used, the probability that a given state S'=i will be observed given S, A and $\omega$ is directly given by:

$$p(S' = i) = \omega p(S' = i|A, S) + (1 - \omega)p(S' = i|S)$$

The probability that the participant predicts the next state would be i (e.g. a square state) when confronted to the hypothetical state-action pair S,A (e.g. circle state, blue action) is finally given by:

$$p(prediction = i) = \frac{\exp(\beta_{choice} p(S' = i)}{\sum_{j=1}^{j=3} \exp(\beta_{choice} p(S' = i))}$$

Where $\beta_{choice} \in [-Inf, Inf]$ determines to which extent the participants will systematically select the most likely transition (i.e. the highest p(S'=i), according to what has been learned) to make their predictions. A very positive $\beta_{choice}$ implies that the participant systematically select this most likely transition. A $\beta_{choice}$ around 0 implies that the participant mostly makes random guesses. And a $\beta_{choice}$ very negative would imply that the participant mostly go against what he/she has learned.

As written above, the key architecture of reference to demonstrate by means of model comparison that participants tracked two sets of transition probabilities and estimated controllability to solve the task is the SAS' model alone. Indeed, such architecture has the same asymptotic performance as the SAS'-SS'-Ω architecture (which thus constitutes a generalization of the SAS' model) in stable environments.

#### 4.8 Model space

Hereafter we describe the 5 different models subjected to model comparison procedure.

1. *SAS' model*. This model only used SA-S' transitions to predict upcoming states. Thus, it had only **2** parameters: a learning rate $\alpha_{sas'}$ and a inverse temperature $\beta_{choice}$. The arbitrator variable $\omega$ was set to a value of 1, so that the spectator model had no impact whatsoever on the prediction processes. No prior injection was used, as it would prevent the actor model from learning in uncontrollable settings. This model had thus **2** parameters.

2. *SAS'-SS'-H*. The update of SA-S' and S-S' transition probabilities was balanced, as the same learning rate was used for the actor and spectator modules (i.e. $\alpha_{sas'} = \alpha_{ss'}$). Counting the two parameters controlling the inference of controllability based on H (i.e. $thres_H$ and $\beta_H$), the inverse temperature parameter $\beta_{choice}$ and the prior injection parameter lambda $\lambda$, this model had thus **5** parameters.

3. *SAS'-SS'-JS*. The update of SA-S' and S-S' transition probabilities was balanced, as the same learning rate was used for the actor and spectator modules (i.e. $\alpha_{sas'} = \alpha_{ss'}$). Counting the two parameters controlling the inference of controllability based on H (i.e. $thres_{JS}$ and $\beta_{JS}$), the inverse temperature parameter $\beta_{choice}$ and the prior injection parameter lambda $\lambda$, this model had thus **5** parameters.

4. *SAS'-SS'-*$\Omega$. This model made use of the full architecture. The update of SA-S' and S-S' transition probabilities was balanced, as the same learning rate was used for the actor and spectator modules (i.e. $\alpha_{sas'} = \alpha_{ss'}$). The update of $\Omega$ was also symmetric, as the learning rate $\alpha_\Omega$ was the same when $\Omega$ increased or decreased. Counting the two parameters controlling the inference of controllability based on $\Omega$ (i.e. $thres_\Omega$ and $\beta_\Omega$), the inverse temperature parameter $\beta_{choice}$ and the prior injection parameter lambda $\lambda$, this model had thus **6** parameters.

We also tested models allowing different learning rates for actor/spectator learning (for models 2-4) or asymmetries in the update of $\Omega$ (for models 4), as well as models which did not allow prior injection (i.e., $\lambda$ fixed at 0.5), resulting initially in a very large model space. However, since none of these models outperformed model 4, we chose to restrict model comparison to this reduced model space for the sake of clarity.

#### 4.9 Model fitting procedure

Model fitting was performed using a Variational Bayesian (VB) estimation procedure using the well-validated VBA toolbox[3]. Compared to non Bayesian methods, this approach has the key advantage of accounting for the uncertainty related to model parameters and hidden states, as well as of informing the optimization algorithm about prior distributions of parameters' values. To the exception of $\beta_{choice}$, parameters were transformed so as to restrict their variation to meaningful intervals. Thus, all learning rates where passed through a sigmoïd function ($\frac{1}{1+e^{-x}}$) limiting their variation to the [0,1] interval. Similarly, the $_\Omega$ parameter was passed through a scaled sigmoid ($-1+\frac{2}{1+e^{-x}}$) so as to limit its variation to the [-1,1] interval. Finally, the $\beta_\Omega$ parameter was constrained to be positive using an exponential transformation.

For the behavioural experiments, the prior distributions of the various learning rates and threshold parameter were innately defined as Gaussian distributions of mean 0 and variance 3, which approximates the uniform distribution over the interval of interest after sigmoid transformation, with a starting value of 0.5. The prior distributions of $\beta_{choice}$ and $\beta_{\omega}$ parameters were defined as Gaussian distribution of mean 0 and variance 10. For the fMRI and the stress experiments, the prior distributions of every parameter was defined using the posterior mean and variance obtained from the 50 participants who passed the behavioural experiment.

Hidden states (i.e values) corresponding to transition probabilities were systematically initialized at 1/3 (equiprobability prior), while Ω was initialized at 0. The VB algorithm was not allowed to update the initial values for hidden states. Contrary to the behavioural experiment, the SS'-SAS' task was split in 4 blocks of equivalent length in the stress and fMRI experiments. Thus, we reinitialized all hidden states at their prior values at the beginning of each block to account for this discontinuity.

#### 4.10 Simulations: model and parameter recovery

In order to ascertain that our task could discriminate participants using the SAS' from those using the best fitting SS'-SAS'-Ω scheme, we simulated 500 participants based on each scheme. For each simulated dataset, parameters were randomly drawn from Gaussian distributions whose means and variances were equal to those observed empirically in the fMRI experiment. Both models were then fitted on the two surrogate datasets of 500 simulated participants using flat priors (mean 0 and variance 3 for all parameters). We then performed one Bayesian group comparisons per dataset, in order to obtain the model selection frequencies of each model.

The quality of parameter recovery for the best-fitting model was estimated by the correlation matrix between the parameters used to generate the simulated data and the recovered parameters.

#### 4.11 Model comparisons

Model comparisons were performed using both fixed-effects and random-effects approaches. Fixed-effects analysis assumes that only one model is used by the whole population and thus sum the information criteria over all participants. For the sake of simplicity, in the main text and figures, we report only the results of the random-effects analyses, which treated model attribution as a random factor potentially model specific using the Bayesian group comparison[4] algorithm of the VBA toolbox.

### 5 Functional Magnetic Resonance Imaging (fMRI)

#### 5.1 fMRI: acquisition

All images were collected using a 3T Siemens Magnetom Prismafit MRI scanner (Erlangen, Germany) with a 32-channel head coil. A T2*-weighted multiband echo planar imaging sequence with acceleration factor 8 (MB8) was used to acquire BOLD-fMRI whole-brain covered images (TR = 700 ms, TE = 39 ms, flip angle = 52, voxel size = 2.4 × 2.4 × 2.4 mm3 , slice gap = 0 mm, and FOV = 210 mm). This state-of-the-art sequencing protocol was optimized from the recommended imaging guidelines of the Human Connectome Project, with the fast acquisition speed facilitating the detection and removal of non-neuronal contributions to BOLD changes (http://protocols.humanconnectome.org/HCP/3T/imaging-protocols.html). The experiment was divided in 4 blocks lasting on average 7.7+/-2.1 minutes (662+/-179 volumes). We recorded participants' heartbeats using the scanner's built-in photoplethysmograph, placed on the right index finger. Respiration was measured with a pneumatic belt positioned at the level of the abdomen. Anatomical images were acquired using a T1-weigted MPRAGE sequence, using a GRAPPA acceleration factor of 2 (TR = 2300ms, TE = 3.03 ms, voxel size = 1x1x1mm, 192 transversal slices, 8° flip angle). Field magnitude and phase maps were also acquired.

#### 5.2 fMRI: preprocessing

fMRI data processing and statistical analyses were performed using statistical parametric mapping (SPM12; Wellcome Trust Centre for Neuroimaging, London, UK). For each session, the first 4 volumes were automatically discarded by the scanner. Functional images were slice-time corrected, unwarped using the field maps and realigned to the mean functional image using a rigid-body registration. Functional images were then coregistered to the anatomical T1. Next, the anatomical image were segmented based on tissue prior probability maps for spatial normalisation employing DARTEL[5] and the resulting normalization matrix was applied to all functional images. Finally, all images were spatially smoothed with a 6mm Gaussian kernel, except in the decoding analysis for which unsmoothed images were used.

#### 5.3 fMRI: mass univariate analysis

Statistical analyses of fMRI signals were performed using a conventional two-levels random-effects approach in SPM12. All general linear models (GLM) described below included the 6 unconvolved motion parameters from the realignment step. We also included the eigenvariate of signals from cerebrospinal fluid (CSF) in our GLM (fourth and lateral ventricular). Moreover, we used a retrospective image correction (RETROICOR) method to regress out physiological noise, using 10 cardiac phase regressors and 10 respiratory phase regressors obtained by expanding cosines and sines of each signal phases to the 5th order. We also included time shifted cardiac rates (lag: +6, +10 and +12s) and respiratory volume (-1 ans +5s) as nuisance regressors.

All regressors of interest were convolved with the canonical hemodynamic response function (HRF). All GLM models included a high-pass filter to remove low-frequency artifacts from the data (cut-off = 96s) as well as a run-specific intercept. Temporal autocorrelation was modeled using an AR(1) process. All motor responses recorded were modeled using a zero-duration Dirac function. We used standard voxel-wise threshold to generate SPM maps ($p<0.001$ uncorrected), unless notified otherwise. All statistical inferences based on whole-brain analyses satisfied the standard multiple comparison threshold ($p(FWE)<0.05$) at the cluster level unless notified otherwise. Prediction error and other parametric regressors were systematically z-scored within block to exclude scaling effects. Reaction time regressors were log-transformed before z-scoring. All statistical inferences based on whole-brain analyses satisfied the standard multiple comparison threshold (p(FWE)<0.05) at the cluster level unless notified otherwise. Cluster-size correction was based on random field theory.

All GLM models included separate onset regressors for motor responses, for prediction trials and for the first trial of each exploratory sequence (where no prediction error was elicited). All models also included parametric regressors for reaction time and $\omega$ (reflecting controllability estimates) on prediction trials. Hereafter, we describe the 6 different variants which were used to generate the results reported in this study:

1. *Independent PE models* (Fig. 4a). In these 2 models, exploratory trials were modeled using one regressor, on top of which decision times and either $\delta_{SS'}$ or $\delta_{SAS'}$ were added as parametric regressors.

2. *PE difference model* (Fig. 4b-c and Fig. S4a). In this model, exploratory trials were modeled using one regressor, on top of which decision times and the difference term $\delta_{ss'} - \delta_{sas'}$ were added as parametric regressors. Because the two controllable rules differed in the frequency of repetition of the same state (very rare in rule C1, common in rule C2), we further included a state repeat regressor, taking the value of 1 whenever a state repeat occurred and 0 otherwise.

3. *Binarized PE model* (Fig. S4b). In this model, we split exploratory trials in four categories based on the prediction errors elicited by the spectator and actor models: trials where both PEs were above their 66th percentile, trials where both PEs were below their 67th percentile, trials where only $\delta_{sas'}$ was above its 66th percentile and trials where only $\delta_{ss'}$ was above its 66th percentile. This procedure was chosen to circumvent collinearity concerns in the estimation of BOLD responses, as the two prediction error terms were positively correlated. On average, 57.4375+/-14.16 trials per participant fell in the “only SS’” and 56.25+/-14.28 trials in “only SAS’” category. An asymmetry in trial distributions with respect to controllability conditions was observed (“only SS’” under controllable rule: 45.28+/-18.87 per participant; “only SAS’” under controllable rule: 19.56+/-8.61 per participant), which reflects the fact that the spectator model was more accurate under uncontrollable conditions whereas the actor model was more accurate under controllable conditions.

4. *Controllability model*. This model was similar to the PE difference model, except that the difference term was replaced by $\delta_{\Omega}$, reflecting controllability update. We also included $abs(\delta_{\Omega})$ in order to control for the overall amount of change in controllability estimates.

5. *Decoding*. For decoding analyses, we modeled each "miniblock" of exploration trials using a separate regressor. Since trials were pooled together, decision times were included as a parametric regressor on motor responses in this model. Note that this model used unsmoothed functional images in order to retain a maximum of information in spatial activation patterns. The searchlight radius was set to 8 millimeters. At the second level, we included a regressor to probe the brain regions in which decoding performances were predicted by the impact of controllability on decision times (z-scored difference in log decision times between

controllable and uncontrollable conditions).

Note that all PE effects were systematically double-checked using the mixed-model approach detailed in the next subsection.

#### 5.4 fMRI: mixed-effects ROI analysis

In order to verify the robustness of our whole-brain results and inspect the time course of our parametric effects of interest, we performed mixed-effects analyses on BOLD signal filtered and adjusted for nuisance regressors. This adjusted signal was extracted from the functional clusters uncovered by whole-brain analyses and segmented into trial epochs from -3 to +16 seconds around the onset of each exploration trial (excluding the first of each streak). We then estimated the effect of each regressor of interest, at each time point, for all subjects simultaneously. Subject identity was included as a random effect and a subject-specific intercept was included. Parametric regressors were z-scored in the same way as in the mass univariate analyses. Importantly, this approach was not used for statistical inference — since doing so would constitute double-dipping — but merely for visualization purposes. The anatomical mask used in Fig. S4c was obtained from[6].

#### 5.5 fMRI: decoding analyses

Decoding analyses were performed using the TDT toolbox[7]. Each mini-block of 6 exploratory trials was arbitrarily coded as +1 (controllable) or -1 (uncontrollable) based on the rule governing transitions. We used a leave-one-run out cross-validation scheme with 100 permutations per subject, so that classes remained balanced for training. Training was performed on the beta values associated with each miniblock (see previous section) using a Support Vector Machine (SVM) classifier (L2-loss function, cost parameter set to 1, Liblinear, version 1.94), without feature selection or feature transformation. Since we did not constrain the testing sets to have balanced classes, balanced accuracies were used when reporting the results of the searchlight analysis (12mm sphere) at the whole-brain level.

### 6 Supplementary Notes

#### 6.1 Link between controllability estimation and exploration

In most cases, estimating environmental controllability requires that the agent explores the space of possible actions as randomly as possible. Conversely, if an agent were to always select the same action $A$ in a given state $S$, the empirical transition probabilities to $S'$, $P(s'|a,s)$ and $p(S'|S)$ would always be equal, independently of the actual logical structure of the environment. From an informational viewpoint, this implies that the agent is *acausal* because $S$ is sufficient to predict $S'$ for an external observer. As a result, controllable and uncontrollable environments are indistinguishable both for the agent and an external observer. More formally, it can be easily proven that $H(S'|S) - H(S'|S,A) > 0$ implies $H(A|S) > 0$:

$$H(S'|S) - H(S'|S,A) > 0$$

By applying Bayes' rule to both conditional entropy terms, we obtain:

$$H(S|S') - H(S) - H(S,A|S') + H(S,A) > 0$$

By applying the chain rule of conditional entropy, we obtain:

$$H(A|S) - H(A|S',S) > 0$$

$$H(A|S) > H(A|S',S)$$

Since the entropy is non-negative by definition, $H(A|S',S) \geq 0$, therefore $H(A|S) > 0$. This relationship implies that some degree of exploration is required to obtain a non-zero transfer entropy (i.e be causal from an information theory perspective) and therefore to estimate controllability (see also, Fig. S1c). In some cases, controllability can however be inferred in the absence of exploration when one leverages prior knowledge about the structure of the ongoing task[8]. For example, in our paradigm, state repetition was only allowed by one of the two controllable rules: knowing this specific feature, participants could in theory distinguish this rule from every other rule by selecting systematically the actions susceptible of producing state repetitions. In practice, our participants only selected these "diagnostic" actions 53+/-8% of the time.

Importantly, the causality measured by transfer entropy is entirely dependent upon the definition chosen to characterize the states and actions $S$, $A$ and $S'$. In this study, the states $S$ and $S'$ were operationally defined as the successive geometrical shapes appearing on the screen, whereas the actions $A$ were defined as the colors selected by participants. This is obviously an abstraction enabling scientific inquiry. The very existence of agents separated from their environment is highly questionable from an epistemological point of view. Instead, the operational meaning of $H(A|S) > 0$ is as simple as "different colors were selected by the participants in front of a given geometrical shape".

Another assumption of the transfer entropy framework is that the decisions of our participants were only susceptible to being conditioned by the geometrical shape displayed on the screen, that is, $S$. Therefore, the fact that transfer entropy increased above zero when participants explored the different options available in controllable contexts should obviously not be equated with the existence of an absolute causality emanating from the agent.

#### 6.2 Link between transfer entropy and causal mediation

The relationship between transfer entropy and causal mediation measures is a complex problem which is still debated by statisticians and information-theorists. Transfer entropy and causal mediation methods are indeed conceptually related, since both can be used under certain conditions to evaluate the existence and the strength of causal relationship within directed acyclic graphs. However, as Ay and Polani have shown[9], the key difference between a simple “observational” transfer entropy measure I(X;Y|W) and the causal measure I(X→Y|W) is that the values taken by X or W should be manipulated through active “interventions” and W kept constant (when estimating direct effects) in the latter case, hence ensuring their independence from the past values of the system or from other variables not included in the model.

Thus, in the context of our study, the key question is to decide whether actions A (i.e. color selection) can be deemed “interventions” by the participants. If this is the case, then the transfer entropy I(S’;A|S) may indeed express the conditional direct effect of actions on future states, I(A→S’|S). However, by associating S to the mediating variable, this analogy entails retrocausality that actions A should in principle be able influence the state S in which they are performed.

Alternatively, following the scheme of Fig. 1a one may treat A as a variable mediating the relationship between S and S', as in I(S→S'|A). However, this latter expression is not appropriate to study controllability. Indeed, a positive I(S→S'|A) simply means that the causal relationship between S and S' does not fully depend on A whereas a zero I(S→S'|A) can be obtained either when S and S' are not causally related or when A fully mediates their relationship (i.e. natural indirect effect), which in turns implies that A should be conditioned on S in the absence of intervention (and therefore not in a proper relation of control with S', as recently discussed[10]). Beyond these considerations, it is also important to note that I(S;S'|A) corresponds to H(S'|A)-H(S'|A,s) which does not provide a spectator model based on which uncontrollable transition could be predicted (although SS' transition probabilities could still be approximated by marginalizing SAS' over actions). It cannot be a priori excluded that the human brain monitors pure AS' transitions rather than SS' transitions, but it is somewhat unlikely that predictions about the consequences of actions are made in complete independence from the context in which they are performed. By contrast, it seems more likely that prediction are made about the transitions between successive states of the environment irrespective of actions, as this latter predictive process can be applied to all situations where one is spectating environmental changes rather than contributing to them.

In sum, while information-theoretic formulations of causal mediation have been recently proposed[11], causal mediation differs from controllability inference in its goal. Indeed, framed as in Fig. 1b, controllability inference aims at determining whether the *unconditioned* behaviour of an agent can modify the evolution of the environment. By contrast, causal mediation is more relevant when one wants to determine whether the *conditioned* behaviour of an agent contributes to the evolution of the environment. In other words, the causal mediation framework relies on an implicit assumption of causal closure whereas controllability inference does not. Finally, transfer entropy can readily be applied to both continuous variables or discrete variables (as in our task), whereas the calculation of causal mediation metrics such as natural direct and indirect effects has been so far mostly applied to continuous variables.

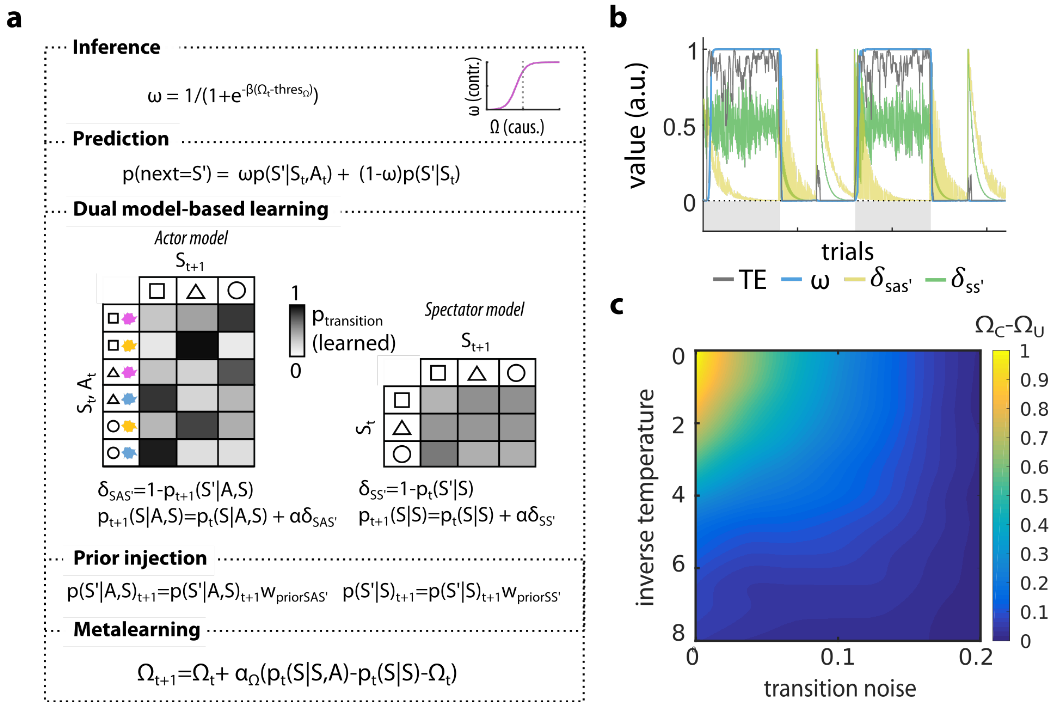

**Figure S1. The SS'-SAS'-Ω architecture. a,** The SS'-SAS'-Ω architecture consists of two parallel probability learning processes. The "actor" model updates on each exploratory trial the transition probabilities from states S and actions A to next states S'. The "spectator" model updates on each exploratory trial the transition probabilities from state S and to next state S': it therefore represents transition probabilities marginalized over actions. These two first-order processes inform a second-order process tracking the expected difference in the probability of observing next states S' given A and S, versus S only. This second order process simply uses a delta-rule to update the variable Ω which reflects the causal influence of one's own action over state transitions. Inferences about controllability are implemented by applying a threshold on Ω determining the value above which it is perceived as evidence that the environment is controllable. This thresholded Ω is further passed into a sigmoid whose slope parameter determines to which extent an Ω below or above the threshold leads to the use of the spectator or the actor model, respectively. The resulting arbitrator variable termed $\omega$ is then used as a weight parameter controlling the mixing of transition probabilities at the time of predictions. Since participants were explicitly presented with the four possible transition rules, they could exploit this knowledge to constrain the update of the first-order models and to some extent, to infer controllability: for example, they knew that state repetitions were only possible under one of the two controllable rules. Prior injections was used to account for this specificity of our task (see Computational modelling section for details). **b,** Time courses of first order prediction errors (green and yellow), transfer entropy (gray, computed within a rolling window of 20 trials) and $\omega$ (blue) in a simulation of the task (noise level=0, learning rates=0.1, $thres_{\Omega} = -0.1$ and $\beta_{\Omega} = 50$, $\lambda = 0.2$). **c,** Discrimination of task conditions (C-U) by Ω in simulations varying transition noise (0 to 0.2) and the inverse temperature parameter determining to which extent the agents choose based on state values (in this simulation, the 3 states gave rewards of 0, 0.5 and 1 units and the agents learned these values using a simple Rescorla-Wagner rule). This shows that Ω keeps discriminating task conditions (C-U>0) even for relatively high noise and conditioning levels.

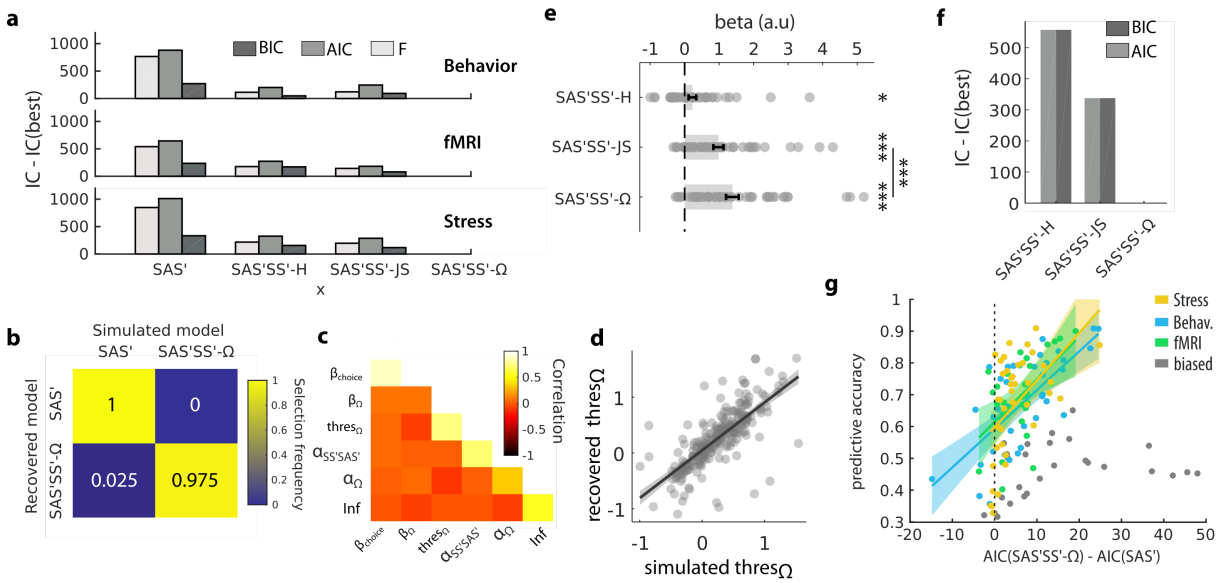

**Figure S2. Advantage of SS'-SAS'-Ω over standard SAS' model-based learning and other controllability estimation schemes. a,** Model comparison treating model attribution as a fixed-effect, using both the AIC and BIC. This analysis split by experiment shows that the SS'-SAS'-Ω architecture outperformed all other candidate models. **b,** Model recovery analysis showed that the task could reliably discriminate the SAS' model alone from the best-fitting SS'-SAS'-Ω scheme. The SAS' model was always identified as the most likely model when it was used to generate the surrogate data, using either BIC or AIC as a goodness of fit metric. When the SS'-SAS'-Ω scheme was used to generate the data, it was identified as the most likely model 97.5% of the time when using BIC (as reported in the panel) and 100% of the time when using AIC. **c,** All parameters of the best-fitting SS'-SAS'-Ω model were recovered correctly with the exception of $thres_{\Omega}$ and to some extent $alpha_{\Omega}$ (r=0.08 and r=0.33, all other r>0.55 in the diagonal). Parameters were not correlated with each other (off-diagonal coefficients between -0.12 and 0.07). **d,** Crucially, the threshold parameter (thresΩ) used to analyse interindividual differences in controllability estimation and to evaluate the effect of the stress induction was highly recoverable (r=0.74). **e,** Logistic regressions attempting to predict whether responses in a pair of prediction pairs would be identical or not based on the value of the arbitrator variable $\omega$ showed that the SS'-SAS'-Ω was more tightly related to variations in this implicit measure of subjective controllability. **f,** Model comparisons corroborated the analysis of beta coefficients reported in e,. **g,** The relative advantage of this best fitting model relative to the standard SAS' model was positively correlated with the mean predictive accuracy across participants in each of the 3 experiments, which demonstrates the benefits of using such arbitrating strategy (behaviour: r=0.58, p<0.001; fMRI: r=0.32, p=0.07; stress: r=0.54, p<0.001). In this analysis, participants who were highly biased towards the spectator model (i.e., belonging to the last quintile of the $thres_{\Omega}$ parameter or $thres_{\Omega} > 0.31$) were treated independently because such bias implied a high reliance on the spectator model (as made possible by the SS'-SAS'-Ω model only) but was typically associated with a low predictive accuracy (0.47+/-0.08 against 0.68+/-0.14 for the rest of the group). Error bars and shaded areas represent SEM.

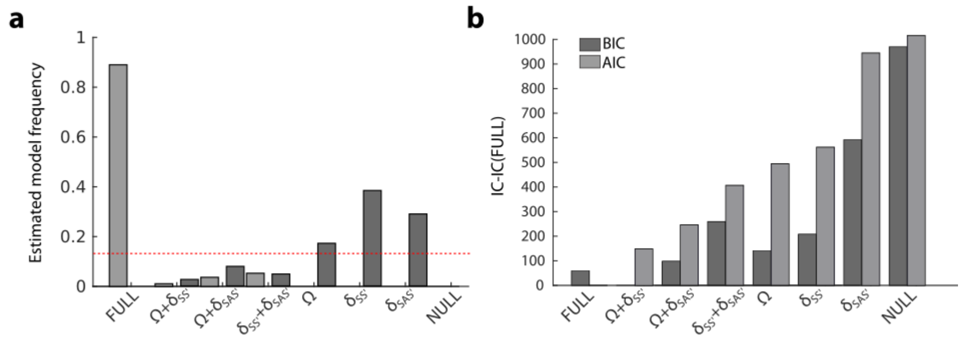

**Figure S3. Model comparison related to the analysis of decision times during exploratory trials. a,** Model comparison treating model attribution as a random effect. When using AIC, the most likely model was the full model which included Ω, $\delta$SAS' and $\delta$SS' as regressors. When using BIC, the most likely model included only $\delta$SS' due to a more stringent penalization for model complexity. As the main goal of this reaction time analysis was to add evidence supporting the dissociation between the actor and spectator model, the fact that the best fitting model based on the BIC criterion included $\delta$SS' constitutes a positive result, given that the existence of actor models is already well-established by a large literature on model-based reinforcement-learning. **b,** Similar conclusions held when treating model attribution as a fixed effect, although the best fitting model based on the BIC also included Ω alongside $\delta$SS' as a predictor in this case.

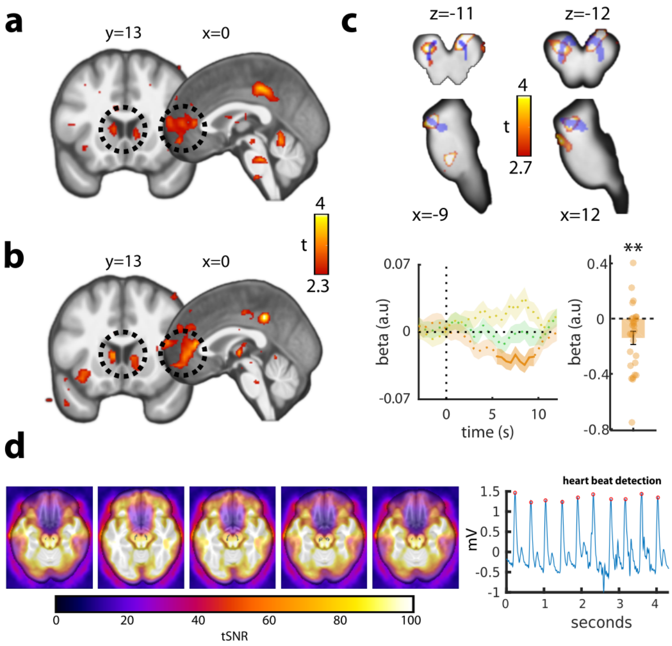

**Figure S4. Supplementary neuroimaging analyses and quality checks. a,** Categorical contrast revealing brain regions where BOLD responses were higher in trials in which only $\delta$SAS' was above its 66th percentile as compared to trials in which only $\delta$SS' was. **b,** Paired t-test contrast opposing the parametric effects of $\delta$SAS' and $\delta$SS' obtained from two separate GLMs. These analyses confirmed the differential encoding of each type of prediction error in the mPFC and to some extent in the nucleus accumbens (see Supplementary Table 3). **c,** Given that the nucleus accumbens and the mPFC are densely irrigated by dopamine which plays a key role in prediction error signaling, we further evaluated BOLD responses in the dopaminergic brainstem. Indeed, a functional cluster overlapping with the substantia nigra (SN) and ventral tegmental area (VTA) could be seen in the maps corresponding to Fig 4b-c and S4a-b. Thus, we analysed brain stem responses of the 27 participants for which respiration and heart rate signals could be used to denoise recordings and improve signal to noise ratio in the VTA (see panel d). Parametric maps (small volume correction: $p < 0.05^{FWE}$) and a ROI analysis within an anatomical mask covering the substantia nigra that the ventral tegmental area [delimited in blue, t(26)=-2.93, p=0.006] confirmed that dopaminergic brain stem encoded the tested for the difference term $\delta$SAS'-$\delta$SS'. The same conclusion was also reached when performing a mixed-effects analysis within the anatomical mask. **d,** The mean tSNR of smoothed normalized data in the dopaminergic brainstem mask (delimited in black at the center of each slice) was comparable to that of ventral cortical structures (mean value of the group: 78.9). Recorded physiological signals allowed reliable detection of individual heart beat and respiration signals, which were used to form retrospective correction of physiological motion (RETROICOR).
Error bars and shaded areas represent SEM.

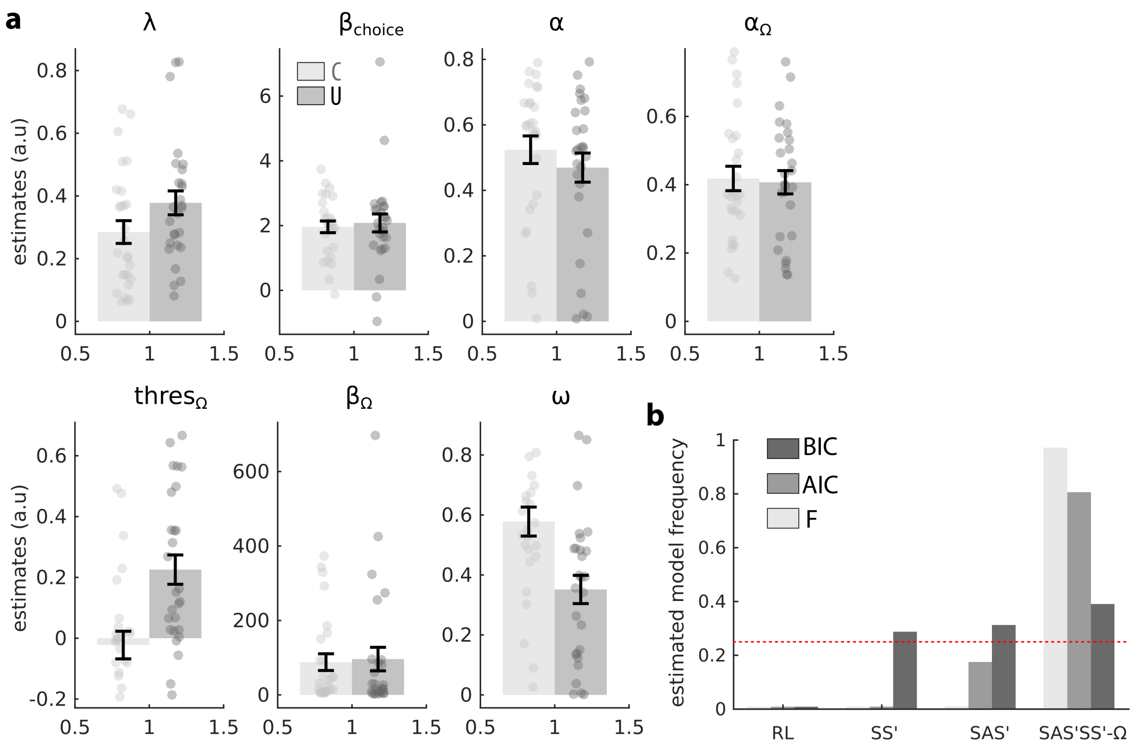

**Figure S5. Additional results related to the stress experiment. a,** Best-fitting parameter and mean value of the arbitrator variable $\omega$ split by stress condition. **b,** Supplementary model comparison restricted to participants who underwent uncontrollable stress. A Bayesian group comparison showed that the SAS'-SS'-Ω was the most prevalent model according to the three information criteria (all exceedance probabilities were above 0.99, except for BIC where Ep=0.58). This model space included a simple value-based RL scheme and a model-based learning scheme exploiting only the spectator model (see Computational modelling section for more details).
Error bars and shaded areas represent SEM.

| | Rule (main effect) | | | Time (main effect) | | | Rule x Time (interaction) | | |
|---|---|---|---|---|---|---|---|---|---|
| | df | F | p | df | F | p | df | F | p |
| behavior | 1 / 47 | 9.58 | 0.003 | 5 / 235 | 4.81 | <0.001 | 5 / 235 | 2.05 | 0.07 |
| fMRI | 1 / 31 | 1.73 | 0.2 | 7 / 217 | 1.94 | 0.06 | 7 / 217 | 1.73 | 0.1 |
| stress | 1 / 53 | 8.29 | 0.006 | 7 / 371 | 2.56 | 0.006 | 7 / 371 | 2.05 | 0.048 |

**Table S1**. Prediction accuracies of participants in the 3 experiments analysed using a 2-way repeated-measures ANOVA. Related to Fig. 2A.

| Label | Abbr. | Side | pFWE | Extent | Peak T | Peak Z | X | Y | Z |
|---|---|---|---|---|---|---|---|---|---|
| Dorsomedial prefrontal | dmPFC | L | <0.001 | 188 | 8.18 | 5.93 | -9 | 11 | 56 |
| Dorsolateral prefrontal | dlPFC | L | 0.036 | 45 | 7.91 | 5.81 | -51 | 2 | 53 |
| Intraparietal sulcus | IPS | R | <0.001 | 226 | 6.91 | 5.34 | 30 | -70 | 38 |
| Dorsolateral prefrontal | dlPFC | R | <0.001 | 238 | 6.88 | 5.32 | 51 | 14 | 38 |
| Anterior Insula | AI | L | 0.004 | 69 | 6.57 | 5.16 | -30 | 26 | -1 |
| Intraparietal sulcus | IPS | L | <0.001 | 134 | 6.40 | 5.07 | -30 | -70 | 38 |
| Dorsolateral prefrontal | dlPFC | L | <0.001 | 137 | 6.01 | 4.86 | -45 | 2 | 32 |
| Anterior Insula | AI | R | 0.044 | 43 | 5.63 | 4.64 | 30 | 26 | -1 |
| Dorsolateral prefrontal | dlPFC | R | 0.001 | 85 | 5.62 | 4.63 | 36 | 5 | 65 |
| Precuneus | | R | 0.011 | 58 | 5.10 | 4.31 | 9 | -67 | 53 |

**Table S2**. Related to Fig. 4a. Clusters associated with the conjunction of parametric regressors $\delta$SS' and $\delta$SAS' (minimal cluster extent: 10, voxel-wise threshold: p<0.001 uncorrected, cluster-wise threshold: p$<$ 0.05$^{FWE}$).

| Label | Abbr. | Side | pFWE | Extent | Peak T | Peak Z | X | Y | Z |
|---|---|---|---|---|---|---|---|---|---|
| **δSS’ - δSAS’ (negative)** | | | | | | | | | |
| Nucleus Accumbens | Nacc | L/R | 0.002 | 76 | 4.74 | 4.08 | -6 | 8 | 5 |
| Medial prefrontal | mPFC | R | 0.004 | 69 | 4.62 | 4 | 9 | 41 | 17 |
| TemporoParietal Junction* | TPJ | R | 0.093 | 34 | 4.02 | 3.58 | 48 | -40 | 35 |
| **(δSAS’>66th & δSS’<67th) > (δSS’>66th & δSAS’<67th)** | | | | | | | | | |
| Medial prefrontal | mPFC | L/R | 125 | 0.015 | 5.02 | 4.26 | -12 | 50 | 14 |
| Posterior cingulate | ACC | L | 336 | <0.001 | 4.16 | 3.68 | -15 | -55 | 35 |
| Nucleus Accumbens* | Nacc | L | 0.99 | 12 | 3.68 | 3.32 | -9 | 11 | 8 |
| TemporoParietal Junction* | TPJ | R | 0.31 | 60 | 3.5 | 3.19 | 48 | -61 | 32 |
| **δSAS’ (parametric) > δSS’ (parametric) - separate GLM** | | | | | | | | | |
| Medial prefrontal (rostral) | rmPFC | R | 102 | 0.046 | 4.23 | 3.73 | 15 | 62 | 5 |
| Medial prefrontal | mPFC | R | 336 | <0.001 | 4.16 | 3.68 | 6 | 44 | 23 |
| Nucleus Accumbens* | Nacc | R | 0.89 | 32 | 3.39 | 3.11 | 9 | 17 | -1 |
| Nucleus Accumbens* | Nacc | L | 0.99 | 14 | 3.86 | 3.46 | -6 | 11 | 5 |
| TemporoParietal Junction* | TPJ | R | 0.88 | 10 | 3.89 | 3.48 | 48 | -43 | 32 |
| **δSAS’ (parametric) > δSS’ (parametric) - same GLM** | | | | | | | | | |
| Medial prefrontal | mPFC | L/R | 225 | <0.001 | 4.24 | 3.76 | 6 | 47 | -1 |
| Nucleus Accumbens* | Nacc | R | 0.47 | 50 | 3.94 | 3.52 | 6 | 5 | 5 |
| Nucleus Accumbens* | Nacc | L | 0.99 | 20 | 4.22 | 3.72 | -6 | 11 | 5 |
| TemporoParietal Junction* | TPJ | L | 0.99 | 17 | 3.24 | 2.99 | 57 | -43 | 38 |

**Table S3**. Related to Fig. 4a and S4a-c. $\delta$sas’- $\delta$ss’: clusters encoding the difference between the two prediction error terms (parametric contrast, minimal cluster extent: 10, voxel-wise threshold: p<0.001 uncorrected, cluster-wise threshold: $p < 0.05^{FWE}$). (SAS’>66th SS’<67th) > (SS’>66th SAS’<67th): clusters responding to the contrast “Only actor PE above the 66th percentile”>”Only spectator PE above the 66th percentile”. SAS’ (parametric) > SS’ (parametric) - separate GLM: clusters where the encoding of $\delta$sas’ was significantly higher than $\delta$ss’ (parametric effects estimated in separate first-level GLMs). SAS’ (parametric) > SS’ (parametric) - same GLM: clusters where the encoding of $\delta$sas’ was significantly higher than $\delta$ss’ (parametric effects estimated in the same first-level GLM). All contrasts: minimal cluster extent: 10; voxel-wise threshold: p<0.005 uncorrected (except $\delta$sas’- $\delta$ss’: p<0.001), cluster-wise threshold: $p < 0.05^{FWE}$. The effects in the Nacc and TPJ marked with an asterisk did not survive for multiple comparisons at the cluster level.

| Label | Abbr. | Side | pFWE | Extent | Peak T | Peak Z | X | Y | Z |
|---|---|---|---|---|---|---|---|---|---|
| Posterior Cingulate | PCC | R | <0.001 | 130 | 5.74 | 4.70 | 6 | -25 | 29 |
| Dorsal Anterior Insula | dAI | R | <0.001 | 98 | 5.32 | 4.45 | 42 | 8 | 17 |
| Medial Prefrontal | mPFC | L | <0.001 | 109 | 5.30 | 4.44 | -6 | 50 | 17 |
| TemporoParietal Junction | TPJ | R | <0.001 | 199 | 5.21 | 4.38 | 60 | -37 | 26 |
| Lateral Occipital | | R | 0.005 | 68 | 4.70 | 4.05 | 42 | -79 | 23 |
| Precuneus | | R | 0.005 | 67 | 4.59 | 3.98 | 21 | -52 | 47 |
| Rostrolateral Prefrontal | | R | 0.037 | 44 | 4.13 | 3.66 | 33 | 50 | 11 |

**Table S4**. Clusters encoding negatively the controllability prediction error term $\delta\Omega$ (minimal cluster extent: 10, voxel-wise threshold: p<0.001 uncorrected, cluster-wise threshold: $p < 0.05^{FWE}$).

| Label | Abbr. | Side | pFWE | Extent | Peak T | Peak Z | X | Y | Z |
|---|---|---|---|---|---|---|---|---|---|
| **Decoding of subjective controllability > balanced accuracy** | | | | | | | | | |
| Dorsolateral prefrontal | dlPFC | R | 0.003 | 272 | 5.63 | 4.61 | 24 | 41 | 32 |
| Dorsolateral prefrontal | dlPFC | L | 0.016 | 202 | 4.89 | 4.16 | -24 | 14 | 44 |
| Precuneus | | R | 0.002 | 296 | 4.51 | 3.91 | 12 | -67 | 41 |
| Supplementary motor | SMA | R | 0.004 | 262 | 4.44 | 3.86 | 69 | -13 | 29 |
| Dorsomedial prefrontal | dmPFC | L | <0.001 | 353 | 4.14 | 3.65 | -3 | 5 | 53 |
| **Decoding of subjective controllability modulated by the effect of controllability on decision times** | | | | | | | | | |
| Dorsolateral prefrontal | dlPFC | R | 226 | 0.009 | 4.76 | 4.08 | 45 | 5 | 29 |
| Cerebellum | | L | 199 | 0.018 | 4.55 | 3.94 | -21 | -49 | -37 |
| Anterior cingulate | ACC | L | 177 | 0.031 | 3.86 | 3.45 | -12 | -1 | 32 |

**Table S5**. Top: Clusters in which the decoding accuracy of upcoming subjective controllability reports exceeded balanced chance levels (p$< 0.05^{FWE}$, voxel-wise threshold: p<0.005 uncorrected). Bottom: Clusters in which the difference in reaction times between controllable and uncontrollable conditions modulated positively the decoding accuracy of upcoming subjective controllability reports (p$< 0.05^{FWE}$, voxel-wise threshold: p<0.005 uncorrected).

| | mean C | std C | mean U | std U | t(26) | p |
|---|---|---|---|---|---|---|
| β | 1.96 | 0.94 | 2.08 | 1.44 | -0.52 | 0.608 |
| β(Ω) | 88.00 | 116.84 | 96.22 | 164.28 | -0.20 | 0.843 |
| thres(Ω) | -0.02 | 0.24 | 0.23 | 0.25 | -4.56 | 0.000 |
| α | 0.52 | 0.22 | 0.47 | 0.23 | 1.06 | 0.299 |
| α(Ω) | 0.42 | 0.19 | 0.41 | 0.18 | 0.24 | 0.810 |
| λ | 0.28 | 0.19 | 0.38 | 0.20 | -2.04 | 0.052 |
| ω | 0.58 | 0.25 | 0.35 | 0.24 | 4.25 | 0.000 |

**Table S6**. Best-fitting parameters estimated in the stress experiment in participants exposed to controllable or uncontrollable stressors. The t- and p-values of independent two-tailed t-tests are reported.
